# Chronology of tRNA structural dynamics prior to and during interaction with a pseudouridine synthase

**DOI:** 10.1101/2025.05.06.652307

**Authors:** Emily M Dennis, Nico E Conoan Nieves, Madison Kadrmas, Abigail L Vaaler, Margaret L Barry, David M Garcia, Julia R Widom

## Abstract

Transfer RNA has long served as an exemplar of a thermodynamically stable, structured RNA. Yet it undergoes significant structural changes upon binding and catalysis by diverse modification enzymes. We leveraged optical binding assays and single-molecule FRET to observe the structural dynamics of two yeast tRNAs, in isolation, and upon interaction with the conserved pseudouridine synthase Pus4/TruB. We show that unmodified and pseudouridylated tRNA^eMet(CAU)^ and tRNA^Thr(AGU)^ all sample open, compact and intermediate conformations, though tRNA^Thr(AGU)^ exhibits faster dynamics. Consistent with its role in modifying RNAs with different structural properties, Pus4 binds robustly to unmodified tRNA, tRNA that was pseudouridylated prior to engaging Pus4 (“pre-modified”), and even an unrelated riboswitch RNA. Pus4 binding to tRNA^eMet(CAU)^ leads to additional tRNA conformational states. The ensemble of conformations explored by Pus4-bound pre-modified tRNA resolved within minutes back into open, compact and intermediate states. tRNA^eMet(CAU)^ that is initially unmodified more gradually approached the ensemble of structures that were attained rapidly by pre-modified tRNA. Thus, Pus4 both catalyzes a lasting chemical change on tRNA and remodels it over time after catalysis, perhaps to promote subsequent steps of tRNA maturation.

## Introduction

Pseudouridine (Ψ) is the most abundant RNA chemical modification in nature. Position 55 in the T-loop is pseudouridylated in nearly all tRNAs, a modification that is most commonly catalyzed by Pus4/TruB/TRUB1+TRUB2 (yeast/bacteria/human). This change from U_55_ to Ψ_55_ creates an additional hydrogen bond donor, increasing base-stacking and helix rigidity, while promoting tertiary interactions between the T-arm and D-arm in the “elbow” region of a folded tRNA. Thus Ψ_55_ contributes to the characteristic three-dimensional structure of tRNA (Lorenz et al. 2017; Zhang and Ferré-D’Amaré 2016; Biela et al. 2023; Robertus et al. 1974; Shi and Moore 2000). In the budding yeast *Saccharomyces cerevisiae*, Pus4 pseudouridylates 41 out of 42 cytosolic tRNA isoacceptors (Shaw et al. 2024), and 23 out of 24 mitochondrial tRNA isoacceptors (Reinsch and Garcia 2025). In *E. coli*, all tRNAs are pseudouridylated by TruB (Cappannini et al. 2024). Therefore, the majority of tRNA molecules in these species are bound and modified by Pus4/TruB, making this enzyme a useful model for understanding the interplay between tRNA structure and dynamics and an enzyme that chemically modifies it.

Previous studies have identified various RNA sequences and/or structures important for binding and modification by various pseudouridine synthase enzymes (Carlile et al. 2019; Lin et al. 2024; Purchal et al. 2022; Behm-Ansmant et al. 2003; Grünberg et al. 2023; Roovers et al. 2006; Gurha et al. 2007; Ansmant et al. 2000, 2001; Zhou et al. 2011; Tillault et al. 2018; Schaening-Burgos et al. 2024; Carlile et al. 2014; Urban et al. 2009; Lecointe et al. 1998; Motorin et al. 1998; Gurha and Gupta 2008), including Pus4/TruB (Mukhopadhyay et al. 2021; Phannachet and Huang 2004; Gu et al. 1998). Yet binding by these enzymes can remodel RNAs in ways distinct from the lasting chemical change that they catalyze. An early study on bacterial TruB noted that growth phenotypes associated with its absence could not be explained by its catalytic activity alone (Gutgsell et al. 2000). Subsequently, a second role was more clearly established: that it aids tRNA folding and functions as a tRNA chaperone (Keffer-Wilkes et al. 2016). Other studies have also demonstrated RNA-chaperone-like roles for other tRNA modifying enzymes (Gc et al. 2020; Johansson and Byström 2002; Copela et al. 2006; Keffer-Wilkes et al. 2020). Thus, to exert their full function, tRNA modifying enzymes should, 1) bind tRNA, 2) catalyze tRNA chemical modification, and, in some cases, 3) remodel tRNA structure. These three components could each independently affect the dynamics of tRNA structure, or do so interdependently. We sought to clarify these activities by addressing the following questions: 1) Do structural ensembles of unmodified tRNA and pseudouridylated tRNA differ? 2) How does binding by Pus4 affect the distribution of tRNA structures over time? 3) Does the outcome depend on whether the tRNA is pseudouridylated prior to engagement with Pus4?

To address these questions, we leveraged single-molecule Förster resonance energy transfer (smFRET) to track the conformational fluctuations of tRNA and biolayer interferometry (BLI) to monitor association of Pus4 to tRNA (**Fig. 1**). We show that Pus4 binds rapidly to both unmodified (U_55_) and pseudouridylated (Ψ_55_, hereafter also referred to as “pre-modified”) tRNAs, and subsequently navigates unmodified tRNA through a pathway of catalysis and remodeling lasting tens of minutes, that is partially bypassed by pre-modified tRNA. This pathway involves a temporary population of conformations not prevalent for either U_55_ or Ψ_55_ tRNA in the absence of enzyme. Thus our results show that a prototypical RNA-modifying enzyme shapes the chronology of tRNA structural dynamics through both its catalysis and remodeling.

**FIGURE 1.**
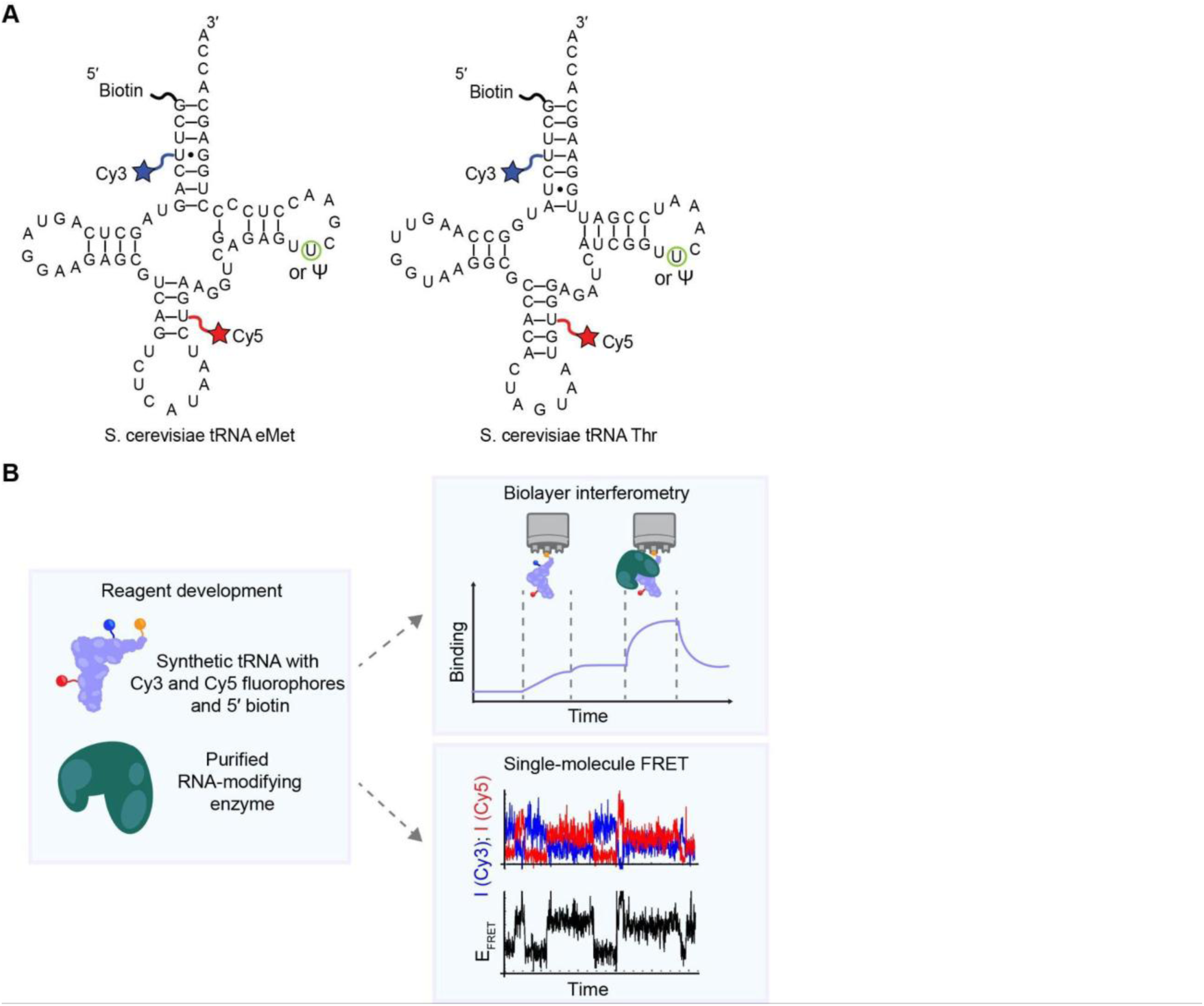
tRNA sequences and methodological approach to investigate how an RNA-modifying enzyme influences tRNA conformation. (*A*) Predicted secondary structures of the tRNA sequences used in this study, including the positions of biotin, Cy3, Cy5 and pseudouridine modification. (*B*) Summary of experimental approach and workflow.

## Results

### Recombinant Pus4 pseudouridylates yeast tRNAs *in vitro*

After establishing a protocol to express and purify GST-tagged, full-length *S. cerevisiae* Pus4 protein from *E. coli* cells (**Supplemental Fig. S1A**), we tested its catalytic activity *in vitro* (**Fig. 2**). Using direct RNA sequencing (DRS), a method we previously established to investigate tRNA modification *in vivo* (Shaw et al. 2024), Ψ_55_ was newly detected on nearly all tRNAs isolated from a *pus4*Δ yeast strain (**Supplemental Table S1**) following a two-minute incubation with an excess of purified Pus4. Modification levels were qualitatively comparable to those previously observed in tRNA isolated from WT strains (Shaw et al. 2024). While some differences in pseudouridine levels were observed between the *in vivo* and *in vitro* data—most notably a lack of modification of tRNAs Arg^ACG^ and Leu^CAA^ *in vitro*—the data supported near-full catalytic activity of purified Pus4 on the vast majority of these cell-derived yeast tRNAs, including the two employed in this study, tRNA^eMet(CAU)^ and tRNA^Thr(AGU)^ (hereafter called tRNA eMet and tRNA Thr, respectively).

**FIGURE 2.**
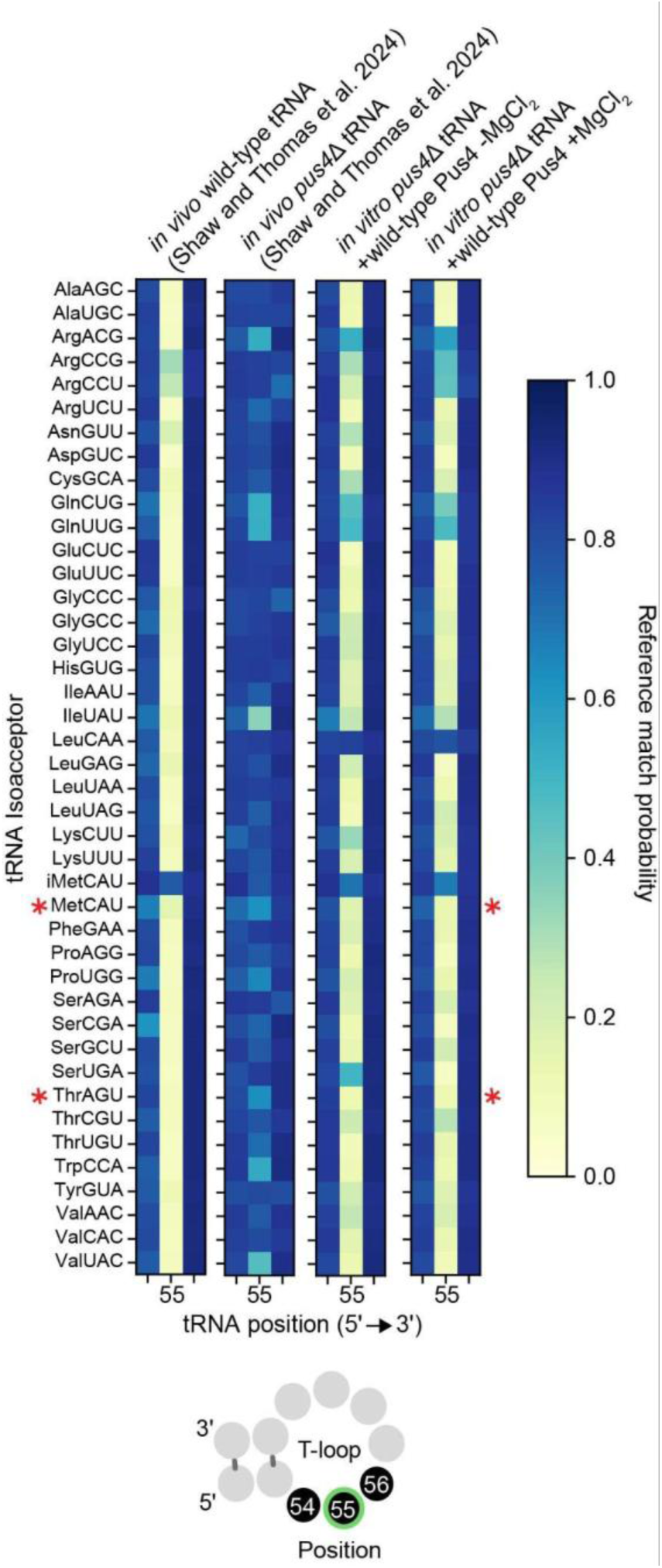
Nanopore direct RNA sequencing shows *in vitro* modification activity of purified Pus4. Heatmaps represent the sequenced nucleotide positions 54–56 of the 42 yeast cytosolic tRNA isoacceptors for tRNA harvested from *pus4*Δ cells and reacted *in vitro* with purified Pus4, with or without 4 mM MgCl_2_. Results from *in vivo* tRNA sequencing are from Shaw et al. 2024. Reference match probability corresponds to the probability that a base-called nucleotide agrees with the reference unmodified nucleotide (A, G, C, or U). More frequently miscalled positions, such as pseudouridine at position 55, produce light yellow squares. When position 55 is base-called as a uridine, the square is dark blue. Red stars indicate the cellular tRNAs corresponding to synthetic tRNAs investigated in this study. At bottom is a diagram of a generic yeast tRNA T-loop with black circles denoting bases shown in heatmaps.

### Synthetic modified tRNAs enable binding and smFRET studies

Having established recombinant Pus4 with desired enzymatic activities, we prepared synthetic tRNA for our biophysical studies of RNA binding by Pus4 (using BLI) and tRNA conformational fluctuations (using smFRET). We prepared synthetic tRNA eMet and tRNA Thr containing either U or Ψ at position 55, with additional modifications required for BLI and smFRET (**Fig. 1A**). Specifically, the 5′ ends were biotinylated for surface immobilization on a BLI probe/microscope slide, the FRET donor fluorophore Cy3 was placed in the acceptor stem, and the acceptor fluorophore Cy5 was placed in the anticodon stem, based on a previous design for human mt-tRNA^Lys^ (Dammertz et al. 2011; Kobitski et al. 2011, 2008; Voigts-Hoffmann et al. 2007).

### Pus4 binds rapidly to tRNA

We utilized BLI to assess the binding specificity of Pus4 and the timescales of its association to and dissociation from both substrate and non-substrate RNAs. RNAs were immobilized on a streptavidin-coated probe that was then passivated to prevent non-specific adsorption of protein, then exposed to buffer containing Pus4 protein (the “association phase”), and finally exposed to buffer lacking protein (the “dissociation phase”) (**Fig. 3A; Supplemental Fig. S2A**). A BLI “shift” is observed when material accumulates on the probe, such as protein binding to immobilized tRNA during the association phase. At a concentration of 200 nM protein, the association curves show rapid binding of Pus4 to tRNA eMet and tRNA Thr, with the accumulation rate slowing after ∼100 seconds (**Fig. 3B,C; Supplemental Fig. S2B,C;** replicate data in **Supplemental Fig. S3**). While accumulation plateaued after ∼15 minutes when Pus4 was present at our highest tested concentration (800 nM), it failed to plateau even after 75 minutes for Pus4 at 50 or 200 nM concentrations (**Fig. 4**). The BLI shift at 200 nM Pus4 eventually converged with the data for 800 nM, suggesting that complete binding occupancy was achieved in this concentration range. We confirmed this using non-denaturing gel electrophoresis as an orthogonal tRNA binding assay, which showed that in free solution, *all* tRNA eMet molecules were bound after 30-minute incubation with Pus4 at concentrations of 200 nM or higher (**Supplemental Fig. S4**). Combined with the prior observation that both tRNA eMet and tRNA Thr were robustly modified within two minutes during the *in vitro* modification experiment (in which Pus4 was present at 1 µM) (**Fig. 2**), we presumed that the majority of the immobilized tRNAs were pseudouridylated in the early period of this association phase once engaged with Pus4.

**FIGURE 3.**
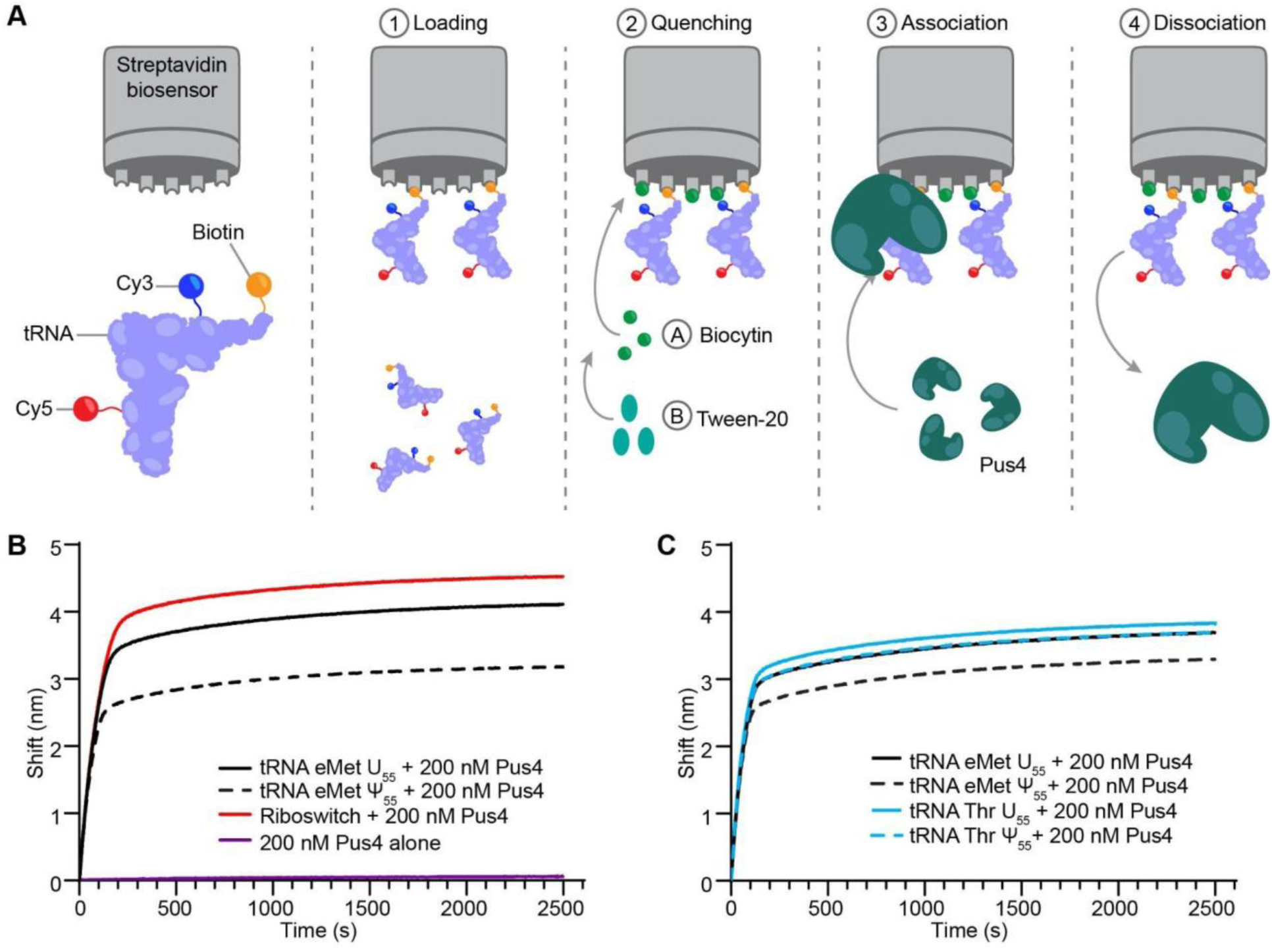
Pus4 binding to tRNA measured using biolayer interferometry. (*A*) Summary of BLI experimental design. Biotinylated and Cy3- and Cy5-labeled tRNAs are immobilized onto streptavidin-coated probes (1) that then undergo two quenching steps and baselining (2). The probe is then transferred into buffer with purified Pus4, where association of Pus4 with immobilized tRNAs occurs (3). The probe is then transferred to a buffer without protein, permitting dissociation (4). (*B*) BLI traces showing the association phase of 200 nM Pus4 to U_55_ (black solid) and Ψ_55_ (black dashed) tRNA eMet, to an unrelated RNA (riboswitch; red), and to a passivated probe lacking RNA (purple). Traces are aligned to start at a shift of 0 nm. (*C*), Same as *B*, but with U_55_ (black solid) and Ψ_55_ (black dashed) tRNA eMet and U_55_ (blue solid) and Ψ_55_ (blue dashed) tRNA Thr. Complete traces in **Supplemental Fig. S2**.

**FIGURE 4.**
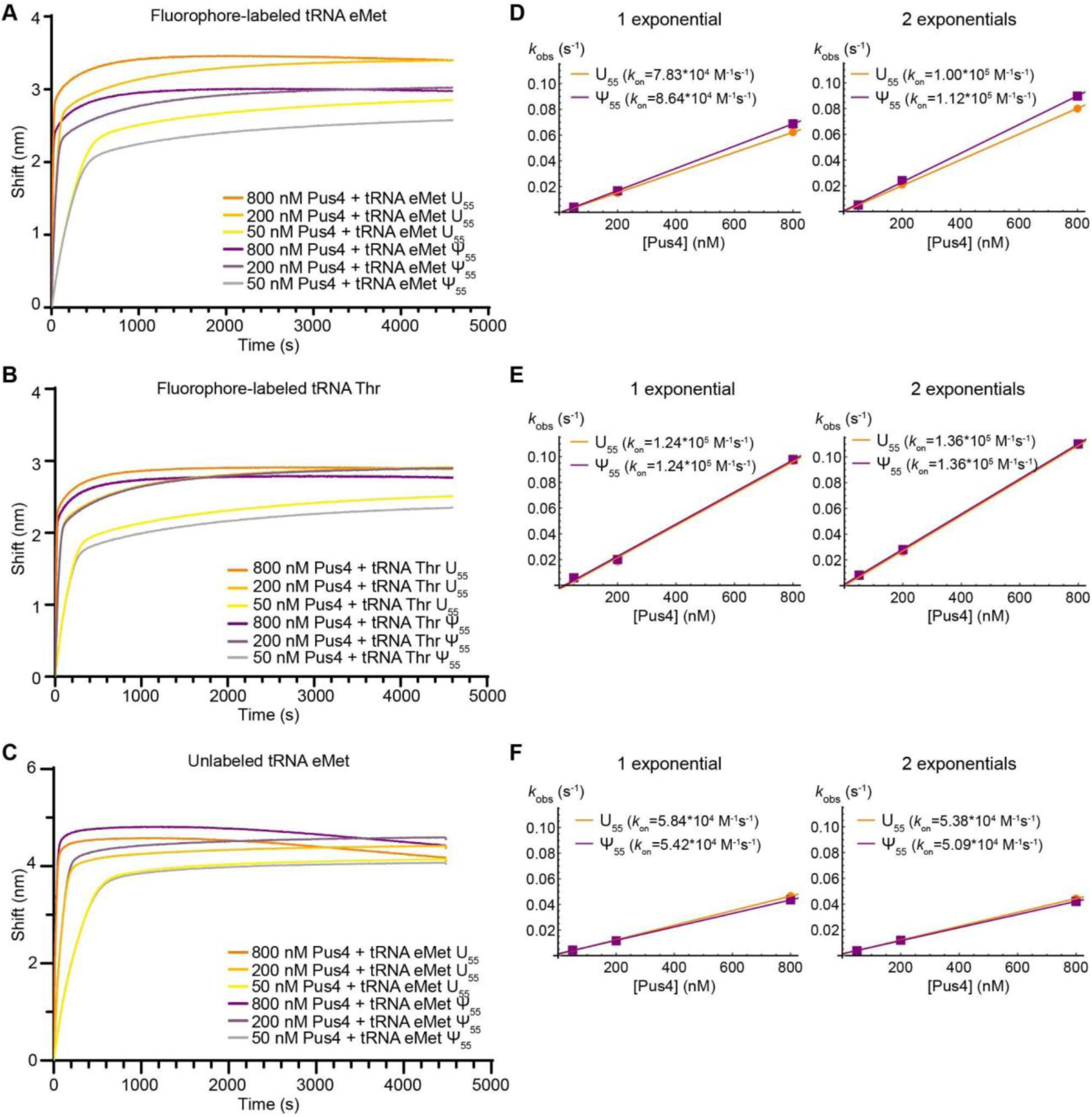
Pus4 binds tRNAs with a high *k_on_*, but a low *k_off_*, with similar affinities for unmodified and pre-modified tRNAs. (*A,B,C*) BLI traces showing the association phase of 50 nM, 200 nM, and 800 nM Pus4 to various tRNAs. Traces are aligned to start at a shift of 0 nm. (*A*) Fluorophore-labeled U_55_ (orange) and Ψ_55_ (purple) tRNA eMet. (*B*) Fluorophore-labeled U_55_ (orange) and Ψ_55_ (purple) tRNA Thr. (*C*) Unlabeled U_55_ (orange) and Ψ_55_ (purple) tRNA eMet. (*D,E,F*), Linear fits to the observed association rate at three Pus4 concentrations. The slope of the fit, representing *kon*, is indicated for each plot. Left columns: *k_obs_* determined from a single exponential fit to the time range of the association phase during which ∼75% of the BLI shift occurs. Right columns: *k_obs_* determined from a double exponential fit to the first 1000 s of the association phase for unlabeled tRNA eMet and the entire association phase for the others. (*D*) Fluorophore-labeled U_55_ (orange) and Ψ_55_ (purple) tRNA eMet. (*E*) Fluorophore-labeled U_55_ (orange) and Ψ_55_ (purple) tRNA Thr. (*F*) Unlabeled U_55_ (orange) and Ψ_55_ (purple) tRNA eMet.

We analyzed the association of Pus4 to tRNA quantitatively by modeling the initial binding of Pus4 as a reversible process occurring under pseudo-first order conditions (details in **Methods**). Mono-exponential fits to the rapid phase of Pus4 association and bi-exponential fits to both phases of association (**Supplemental Fig. S5**) both revealed that the on-rate (*k_on_*) was slightly faster for tRNA Thr (1.36*10^5^ M^-1^s^-1^ from the bi-exponential fit) than for tRNA eMet (1.00*10^5^ M^-1^s^-1^) (**Fig. 4D, E**). These on-rates are comparable to ones previously observed for yeast Pus1 binding to tRNA^Phe^ and tRNA^Val^ (4-6*10^5^ M^-1^s^-1^) (Arluison et al. 1999). These fits could in principle have been used to obtain *k_off_*, however the off-rate during the initial phase of Pus4 accumulation was indistinguishable from zero. When the BLI probe was transferred into a solution lacking protein, an initial fast phase of dissociation was observed, but even after 75 minutes, less than 10% of the shift observed in the association phase was reversed (**Supplemental Fig. S2B,C**). This suggests that Pus4 binds all RNAs investigated with high affinity and dissociates very slowly.

To test whether the fluorophores altered Pus4 binding affinity to tRNAs, we prepared a synthetic tRNA eMet with the same sequence as above, but without Cy3 and Cy5 fluorophores (**Fig. 4C; Supplemental Fig. S6**). The BLI curves showed the same overall shape with a somewhat lower *k_on_* value (5.84*10^4^ M^-1^s^-1^) than observed for labeled tRNA eMet, an impact comparable in size to the variation between different tRNAs (labeled tRNA eMet vs. labeled tRNA Thr). We conclude that while the labels exert a small effect on Pus4 binding, in general our strategy for investigating the conformational ensemble of tRNA using smFRET is not confounded by the effects of these fluorophores on the binding behavior of Pus4.

After the initial binding phase that was used for kinetic analysis, we noted other features of Pus4 binding behavior. In the highest concentration tested (800 nM protein), we observed that after binding plateaued, a mild dissociation occurred slowly over the remainder of the measurement (**Fig. 4**). The effect was more distinct in the unlabeled tRNAs in comparison to the fluorophore-labeled, but was present in all tRNAs. This appeared to be a distinct phase from the binding phase; thus we did not incorporate it into our kinetic analysis. Secondly, the slow, steadily increasing second phase of the association curve observed at all concentrations of Pus4 tested may represent 1) gradual “trapping” of tRNA or Pus4 molecules that began the association phase in conformations not amenable to binding, or 2) Pus4 multimerization such that the stoichiometry of Pus4:tRNA exceeded 1:1 on some of the immobilized tRNA molecules. Multimerization is supported by the shift of Pus4:tRNA bands to the wells for the highest Pus4 concentrations (**Supplemental Fig. S4**), and might be facilitated by the GST tag that is known to dimerize (Fabrini et al. 2009). Our attempts to prepare untagged Pus4, however, were unsuccessful as it showed a strong propensity to aggregate after tag cleavage. To measure the extent to which Pus4 multimerization could impact tRNA binding, we incubated total tRNA isolated from yeast cells with recombinant Pus4, and performed sucrose gradient ultracentrifugation to separate protein:RNA complexes (**Supplemental Fig. S7**). If multimerization of Pus4 under these conditions reduced tRNA binding, we would expect no evident shift in the distribution of protein complexes upon addition of tRNA. However, addition of tRNA caused a global shift of Pus4 to fractions in the gradient containing a higher percentage of sucrose, forming complexes with an apparent molecular weight of approximately 200 kDa (implying complexes of two GST-Pus4 with two tRNAs). Additionally, our smFRET measurements (described below) were performed on sparsely surface-immobilized tRNAs, in which the average distance between tRNA molecules greatly exceeds the size of a protein dimer or even multimer. This would preclude the formation of complexes containing more than one tRNA molecule. In summary, while we can detect Pus4 multimers in our experimental conditions, we do not believe that they represent the dominant form of Pus4 that is interacting with immobilized tRNAs that are the principle substrate that we measured, and moreover our Pus4 samples demonstrated robust catalysis (**Fig. 2**).

### Pus4 also binds robustly to pre-modified tRNA and non-substrate RNA

To test whether Ψ_55_ altered the binding of Pus4 to tRNA, we tested synthetic tRNAs that had pseudouridine present at position 55 using BLI. Although the BLI curve of Ψ_55_ tRNA eMet exhibited a smaller maximum shift than that of U_55_ (**Fig. 4A**), non-denaturing gel electrophoresis indicated that this still corresponded to complete binding occupancy at Pus4 concentrations at or above 200 nM (**Supplemental Fig. S4**). Furthermore, the overall association and dissociation profile was similar between the unmodified and pre-modified variants of both tRNAs, and *k_on_* values were unchanged or slightly higher with Ψ_55_ tRNA (1.12*10^5^ M^-1^s^-1^ for Ψ_55_ tRNA eMet and 1.36*10^5^ M^-1^s^-1^ for Ψ_55_ tRNA Thr), indicating weak discrimination between binding substrate and product. We also performed experiments with an RNA that is not a natural substrate of Pus4 but is similarly sized to tRNA: the bacterial class II NAD^+^ riboswitch, a 61 nt RNA that adopts a pseudoknot structural motif to regulate translation initiation (Conoan Nieves and Widom 2024; Panchapakesan et al. 2021). As we observed for tRNA, Pus4 rapidly associated with the riboswitch, followed by slow continued accumulation and slow dissociation (**Fig. 3B, Supplemental Fig. S2B**). It is worth noting that previous studies have shown pseudouridine synthases can be promiscuous binders of RNA (Carlile et al. 2019; Purchal et al. 2022; Behm-Ansmant et al. 2003; Grünberg et al. 2023), consistent with their roles of modifying multiple sequences and structures in various classes and sizes of RNA. For example, Pus1 has been found to have broad affinity for RNA, even those lacking defining tRNA structural features, as well as tight binding to catalytic products (e.g., K_D_ = 14 nM for pseudouridylated tRNA^Val^) (Arluison et al. 1999). In addition to tRNA, Pus4 modifies mRNA (Carlile et al. 2014; Lovejoy et al. 2014; Schwartz et al. 2014) and mt-rRNA (Begik et al. 2021), *in vivo*. Our data suggest that, like Pus1, Pus4 also exhibits promiscuous binding. This may promote Pus4’s ability to facilitate RNA remodeling more generally, even if its primary catalytic substrate is tRNA.

### tRNA undergoes dynamic fluctuations between three predominant conformational states

We utilized smFRET to investigate how tRNA conformations are affected by Pus4 binding and pseudouridylation. The biotinylated- and fluorophore-labeled tRNA constructs used for BLI were immobilized on microscope slides, and fluorescence from single tRNA molecules was recorded using wide-field prism-based total internal reflection fluorescence (TIRF) microscopy. We recorded short movies (∼2 minutes) sequentially across different regions of the slide of tRNA alone, as well as tRNA during up to 30 minutes of incubation with 200 nM Pus4 protein. Cy3 and Cy5 fluorescence intensity traces were extracted from individual molecules, and data were combined to achieve ∼200 traces for each condition. In order to observe the evolution of the conformational landscape over tens of minutes, the data were combined into discrete time bins of 2–6 minutes, 7–9 minutes, 10–15 minutes and 30–34 minutes after introduction of the imaging buffer, either with or without Pus4. FRET efficiency (E_FRET_) histograms were compiled from the first 50 frames of all traces within each condition.

To accurately recover signals reflecting the distance between the donor and acceptor fluorophores, and therefore the structure of the tRNA, FRET traces were corrected for 1) crosstalk between detection channels, and 2) gamma (γ) factor, which quantifies the relative detection efficiencies and fluorescence quantum yields of Cy3 and Cy5 (**Supplemental Fig. S8**). We determined the γ factor utilizing changes in donor and acceptor intensity upon acceptor photobleaching (McCann et al. 2010), which yielded a distribution with a median value of 2.4 in the absence of Pus4 and 3.0 in the presence of Pus4 (**Supplemental Fig. S8C,D**). Given the significant adjustment to the raw data imposed by a γ factor so far from 1 (**Supplemental Fig. S8A**), we also tested whether utilizing a molecule-by-molecule γ factor correction would improve the resolution of different FRET states. However, restricting the analysis to molecules for which individual γ values could be calculated (molecules in which the acceptor photobleached first) skewed the histogram without sharpening its features (**Supplemental Fig. S8E-G**). We thus elected to utilize the median of the distribution (with or without Pus4 as appropriate) as a global γ correction factor, permitting inclusion of all traces.

Inspection of the corrected traces revealed that in the absence of protein, all four tRNA variants investigated—U_55_ and Ψ_55_ tRNA eMet and U_55_ and Ψ_55_ tRNA Thr—dynamically sampled at least three distinct conformational states (**Fig. 5A-D**). The shapes of the distributions and FRET efficiencies, however, differed between the eMet and Thr tRNA sequences. In the histograms of U_55_ and Ψ_55_ tRNA eMet, which were well-fit with three Gaussian distributions, a low-FRET efficiency feature (LF)—representing a conformation in which the fluorophores are most separated from each other relative to other states—is centered around E_FRET_ = 0.1. An LF feature could arise from molecules that are either less structured or that adopt a structure such as an extended helix that enforces separation of the fluorophores. Mid-FRET (MF) and high-FRET (HF) features—representing conformations in which the fluorophores are closer to each other relative to LF—were observed for both U_55_ and Ψ_55_ tRNA eMet and centered around E_FRET_ = 0.48 and 0.83, respectively (**Fig. 5E,F**). The HF state was moderately more populated in U_55_ than Ψ_55_ tRNA eMet. LF, MF and HF states were previously observed in smFRET analysis of the human mt-tRNA^Lys^ and were assigned to unfolded (U), extended hairpin (E), and cloverleaf-based-L-shape (C) conformations, respectively (Dammertz et al. 2011). The histograms of U_55_ and Ψ_55_ tRNA Thr exhibited broad, asymmetric distributions centered around E_FRET_ = 0.6. These were well-fit with either four (U_55_) or three (Ψ_55_) Gaussian distributions (**Fig. 5G,H**) with centers ranging from E_FRET_ = 0.38 to 0.99, consistent with the variety of states observed in the traces (**Fig. 5C,D**).

**FIGURE 5.**
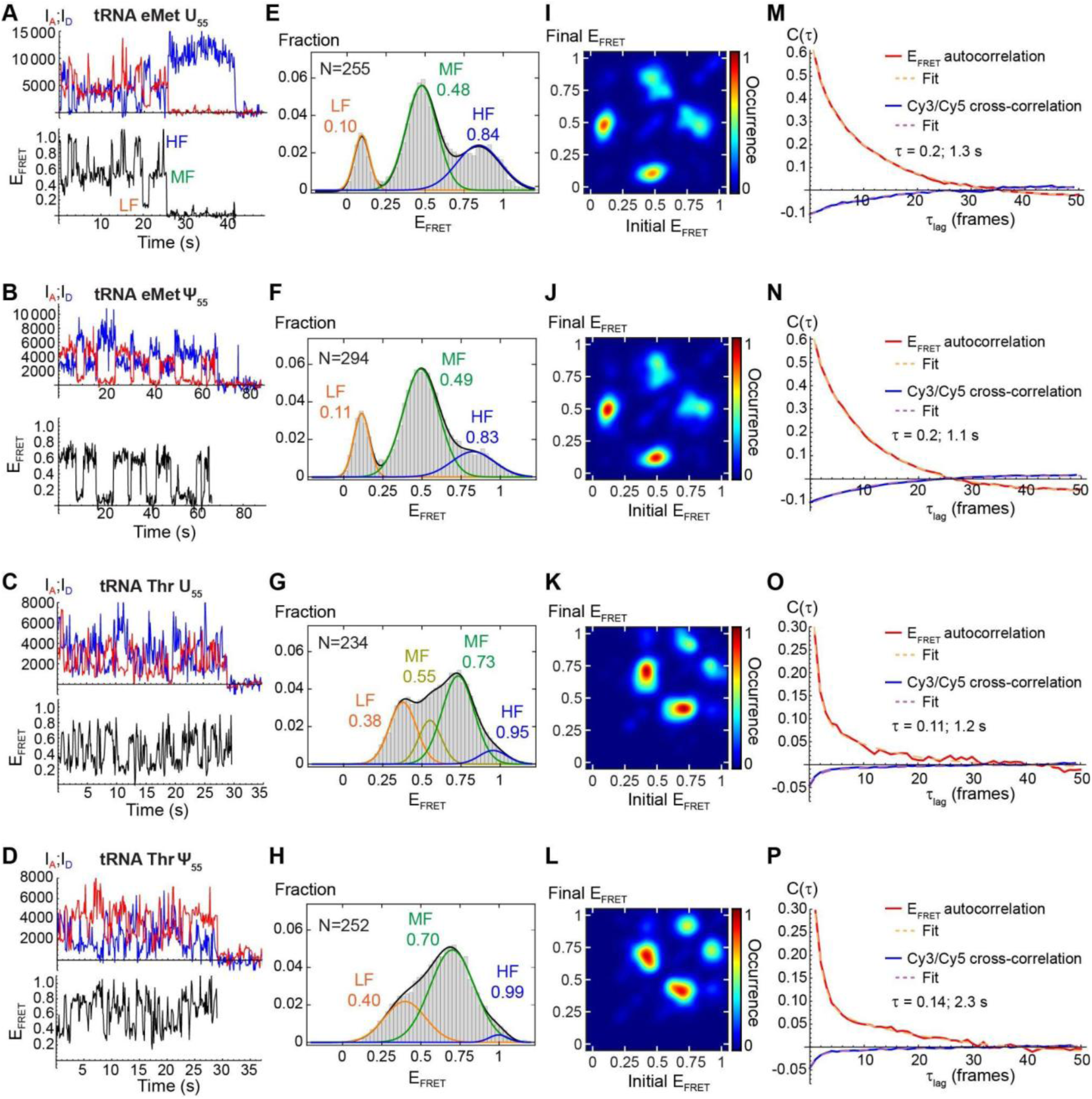
smFRET data on tRNAs in the absence of Pus4. (*A*–*D*) Representative smFRET traces of (*A*) U_55_ tRNA eMet; (*B*) Ψ_55_ tRNA eMet; (*C*) U_55_ tRNA Thr; (*D*) Ψ_55_ tRNA Thr. Top: Donor (blue) and acceptor (red) intensities. Bottom: FRET efficiency. Occurrences of the LF, MF and HF states are indicated in panel *A*. For clarity, the E_FRET_ traces are truncated after donor photobleaching. Only trace segments prior to photobleaching of either dye were subjected to further analysis. (*E*–*H*) E_FRET_ histograms compiled from the first 50 frames of the number of traces indicated by “N” for (*E*) U_55_ tRNA eMet; (*F*) Ψ_55_ tRNA eMet; (*G*) U_55_ tRNA Thr; (*H*) Ψ_55_ tRNA Thr. 3- or 4-Gaussian fits are overlaid with the peak centers labeled. (*I*–*L*) TODPs for (*I*) U_55_ tRNA eMet; (*J*) Ψ_55_ tRNA eMet; (*K*) U_55_ tRNA Thr; (*L*) Ψ_55_ tRNA Thr. The intensity of the heatmap at position (X,Y) quantifies the fraction of molecules that transition from E_FRET_ = X to E_FRET_ = Y at least once during the measurement. (*M*–*P*) FRET autocorrelation (red) and Cy3/Cy5 cross-correlation (blue) functions for (*M*) U_55_ tRNA eMet; (*N*) Ψ_55_ tRNA eMet; (*O*) U_55_ tRNA Thr; (*P*) Ψ_55_ tRNA Thr. Global bi-exponential fits with the same time constants for autocorrelation and cross-correlation functions are overlaid in dashed lines, and the two time constants are indicated.

We analyzed the dynamics of the four tRNAs by performing Hidden Markov Modeling (HMM) (Blanco and Walter 2010). For tRNA eMet, we used a 4-state model (**Supplemental Fig. S9A**) that included the LF, MF, and HF states, in addition to a second HF state (HF2) centered at E_FRET_ = 0.95, which was observed in a fraction of molecules too small to yield a distinct feature in the histogram (5% in U_55_ and 2% in Ψ_55_). The model allowed for transitions between all pairs of the LF, MF and HF states, with the HF2 state occupied only statically, in accordance with the small number of traces that exhibited it. The isolated HF2 state was omitted for tRNA Thr based on inspection of traces, and the E_FRET_ values of the LF, MF and HF states were adjusted to match those obtained from 3-Gaussian fits to their histograms (**Supplemental Fig. S9B**). The pathways of transitions among these states were visualized using transition occupancy density plots (TODPs), which show the prevalence of dynamic or static behavior across hundreds of molecules within each condition (**Fig. 5I-L**). A feature on the diagonal axis of a TODP (i.e. lower left corner to upper right corner) indicates a population of static molecules that remain in one state throughout their measurement, while an off-diagonal feature at position (X,Y) indicates a population of dynamic molecules that transition from states with E_FRET_ = X (initial) to E_FRET_ = Y (final) at least once during the measurement. This analysis showed that the majority of molecules (>80% for U_55_ and Ψ_55_ variants of both tRNAs) exhibited dynamic behavior involving fluctuations between conformations. This is consistent with previous bulk (Cole et al. 1972; Bhaskaran et al. 2014) and smFRET (Voigts-Hoffmann et al. 2007) studies that showed that tRNAs can interconvert between multiple secondary and tertiary structures at physiological salt and ambient temperature. In U_55_ and Ψ_55_ variants of both tRNAs, bright off-diagonal features in the TODPs indicate that molecules readily and reversibly interconverted between the LF and MF states and between the MF and HF states (**Fig. 5I-L**). Significantly, almost no intensity was observed at the locations corresponding to transitions between the LF and HF states in any of the four tRNAs investigated, indicating that interconversion between these two states requires a molecule to transition through an intermediate MF state.

For U_55_ and Ψ_55_ tRNA eMet, HMM analysis further revealed transition rates of 0.3–0.8 s^-1^ between LF and MF states, and slightly faster transition with rates of 0.8–1.5 s^-1^ between MF and HF states (**Table 1**, **Supplemental Fig. S10**). The overall rates of conformational fluctuations in each tRNA were also quantified in a model-free manner using cross-correlation and auto-correlation analyses (details in **Methods**), which identify how rapidly Cy3 and Cy5 intensities become uncorrelated with one another within a collection of smFRET traces. Consistent with the HMM results, the correlation analysis confirmed that fluctuations in FRET efficiency occurred on a time-scale of ∼1 s in U_55_ and Ψ_55_ tRNA eMet (**Fig. 5M,N**). For U_55_ and Ψ_55_ tRNA Thr, fluctuations were notably faster, with HMM-derived transition rates between the LF and MF states almost 4x faster than observed for tRNA eMet (**Table 1**), and biphasic correlation functions dominated by fluctuations on a time-scale <0.15 s rather than the ∼1 s observed for tRNA eMet (**Fig. 5O,P**). This approaches the 100 ms time resolution of our smFRET measurements, suggesting that the indistinct features of the histograms of tRNA Thr may have arisen in part from time averaging across transitions between states. In summary, our analysis revealed that U_55_ and Ψ_55_ tRNA eMet and tRNA Thr all undergo frequent conformational changes involving substantial adjustments in their three-dimensional architecture, with their MF states acting as near-obligate intermediates between LF and HF states.

**TABLE 1.** HMM-derived rate constants (weighted average from a bi-exponential fit) for transitions between each pair of interconverting FRET states. Fits shown in **Supplemental Fig. S10.**

| tRNA | $k_{LF \rightarrow MF} \text{ (s}^{-1}\text{)}$ | $k_{MF \rightarrow LF} \text{ (s}^{-1}\text{)}$ | $k_{MF \rightarrow HF} \text{ (s}^{-1}\text{)}$ | $k_{HF \rightarrow MF} \text{ (s}^{-1}\text{)}$ |
| --- | --- | --- | --- | --- |
| U<sub>55</sub> tRNA eMet | 0.58 | 0.34 | 1.00 | 1.43 |
| Ψ<sub>55</sub> tRNA eMet | 0.79 | 0.58 | 0.79 | 1.49 |
| U<sub>55</sub> tRNA Thr | 2.83 | 1.26 | 0.90 | 1.59 |
| Ψ<sub>55</sub> tRNA Thr | 2.64 | 1.22 | 0.91 | 1.70 |

### Pus4 alters the conformational states and transition dynamics of tRNA eMet

After Pus4 was introduced into a sample chamber containing immobilized U_55_ or Ψ_55_ tRNA eMet, a rapid increase in brightness of both Cy3 and Cy5 indicated that the protein was binding to the RNA (**Supplemental Table S2**), a phenomenon known as “protein-induced fluorescence enhancement” (PIFE) (Hwang et al. 2011). This increase was largely complete by the earliest time window that we probed by smFRET, 2–6 minutes, which begins when the initial fast phase of Pus4 association observed by BLI is nearly complete, and continues into the slow accumulation phase (**Fig. 3B,C**). Concurrently, an increase in conformational complexity was observed, indicating that the protein changed the free energy landscape of tRNA molecules. 2–6 minutes after addition of protein, the histogram of U_55_ tRNA broadens and all three features shift to higher E_FRET_ (**Fig. 6A**, **Fig. 7A**), potentially indicating changes to specific tertiary contacts previously held. Inspection of traces suggested that the observed increase in complexity originates from interconversion between mid-FRET “microstates” with E_FRET_ ∼0.4 and 0.7 (**Supplemental Fig. S11A**). Although we used only one MF state in the HMM model to avoid overfitting and facilitate comparison to data recorded in the absence of protein (**Supplemental Fig. S9A**), the presence of additional microstates is reflected in TODPs. Specifically, the features present in tRNA alone became broader in the presence of Pus4, and transitions were observed between states with a greater variety of E_FRET_ values (**Fig. 5I,J** vs. **Fig. 8A**). We hypothesize that these microstates involve remodeling of the T-loop in order to facilitate pseudouridylation, as observed in crystallography data of the bacterial homolog TruB with tRNA (Pan et al. 2003; Hoang and Ferré-D’Amaré 2001). The shift to higher E_FRET_ in the histogram is partially reversed in the 7–9 minute time window (**Fig. 6**). The non-monotonic changes to the E_FRET_ distribution over time, despite protein occupancy increasing monotonically throughout, (**Fig. 3**), suggests that the protein-RNA complex evolves in structure after its initial assembly. By 30–34 minutes after introduction of Pus4 (at which point our gel shift assay indicated that all tRNA molecules were bound by Pus4 in solution; **Supplemental Fig. S4**), the conformational ensemble resolves back into distinct LF, MF and HF features, with MF and HF centered at higher E_FRET_ than before exposure to Pus4 (**Fig. 7A**). While our approach precluded real-time, simultaneous measurement of pseudouridylation of the immobilized tRNA, our *in vitro* pseudouridylation assay showed that the majority of tRNAs were pseudouridylated by Pus4 within 2 minutes (**Fig. 2**). Therefore, results over the course of these smFRET experiments reflect the enduring influence of Pus4 binding and catalysis on the trajectory of the tRNA conformations sampled.

**FIGURE 6.**
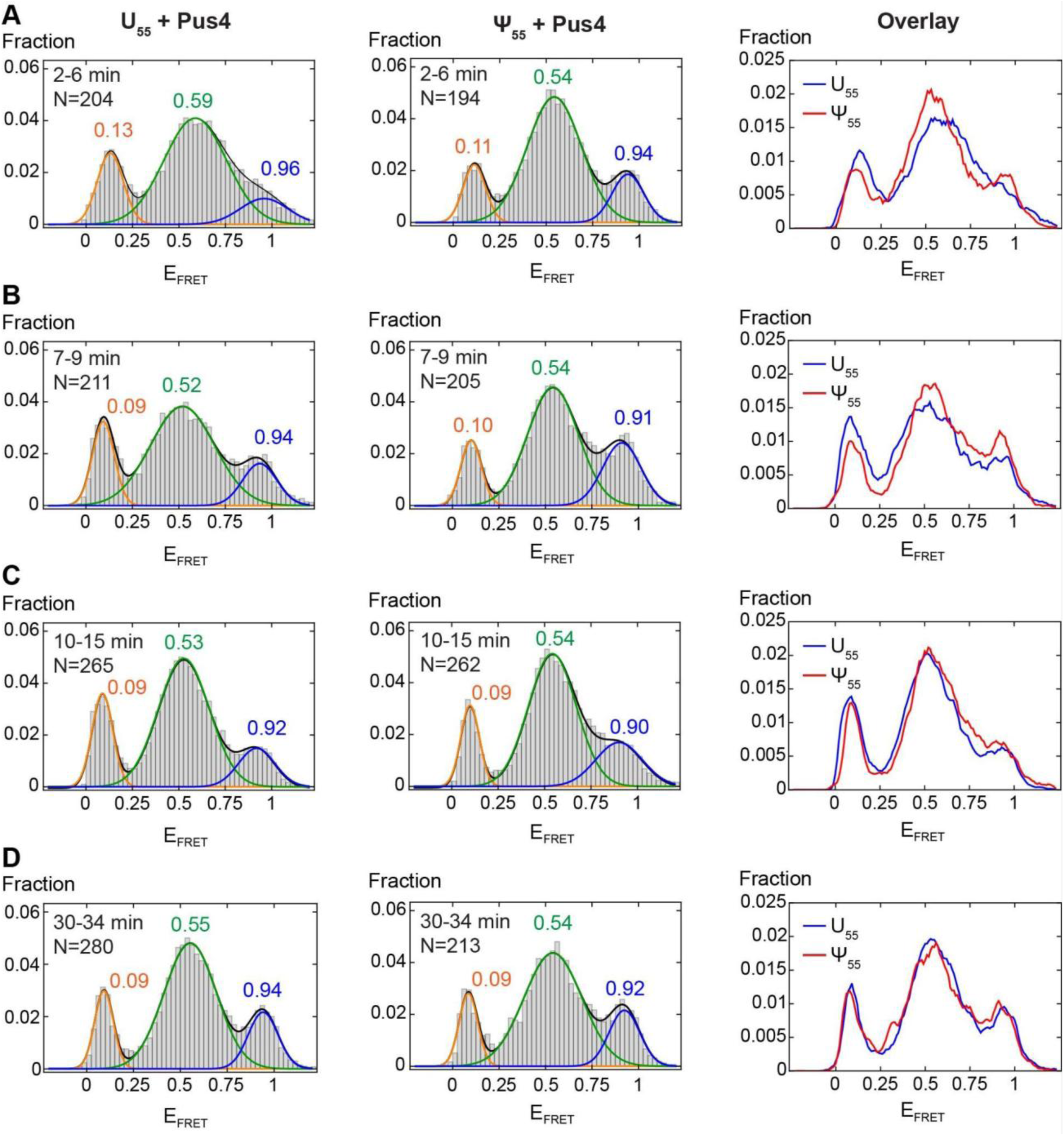
smFRET histograms for U_55_ and Ψ_55_ tRNA eMet in the presence of 200 nM Pus4. (*A*) Histograms compiled from movies recorded 2–6 minutes after injection of Pus4. Left column: U_55_+Pus4 histogram with 3-Gaussian fit overlaid; Middle column: Ψ55+Pus4 histogram with 3-Gaussian fit overlaid; Right column: Overlay of U_55_ and Ψ_55_ histograms. The E_FRET_ value of each state is indicated. (*B*) Same as *A* but compiled from movies recorded 7–9 minutes after injection of Pus4. (*C*) Same as (*A*–*B*) but compiled from movies recorded 10–15 minutes after injection of Pus4. (*D*) Same as (*A*–*C*) but compiled from movies recorded 30–34 minutes after injection of Pus4.

**FIGURE 7.**
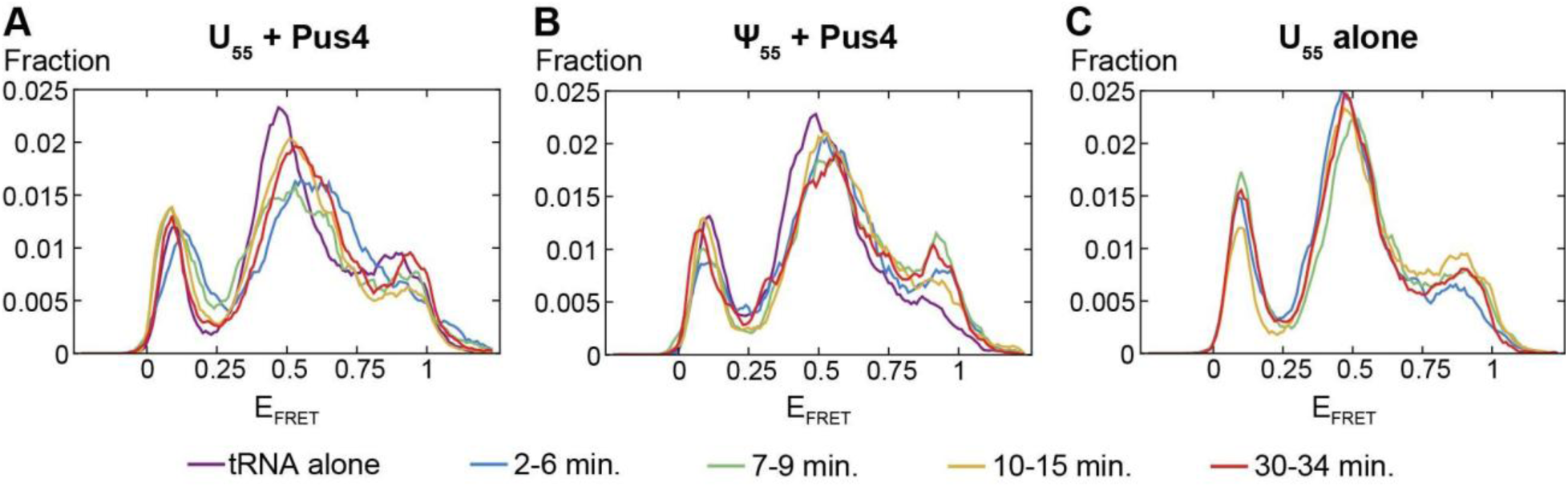
Overlaid smFRET histograms throughout incubation of (*A*) U_55_ tRNA eMet with 200 nM Pus4, (*B*) Ψ_55_ tRNA eMet with 200 nM Pus4, (*C*) U_55_ tRNA eMet with imaging buffer lacking Pus4. Blue: Histogram compiled from traces recorded 2–6 minutes after introduction of imaging buffer containing or lacking Pus4.; Green: 7–9 min.; Yellow: 10–15 min.; Red: 30–34 min. Panels *A* and *B* additionally show the histograms of the corresponding tRNA alone after 10–15 minutes of incubation with imaging buffer lacking Pus4 (purple).

**FIGURE 8.**
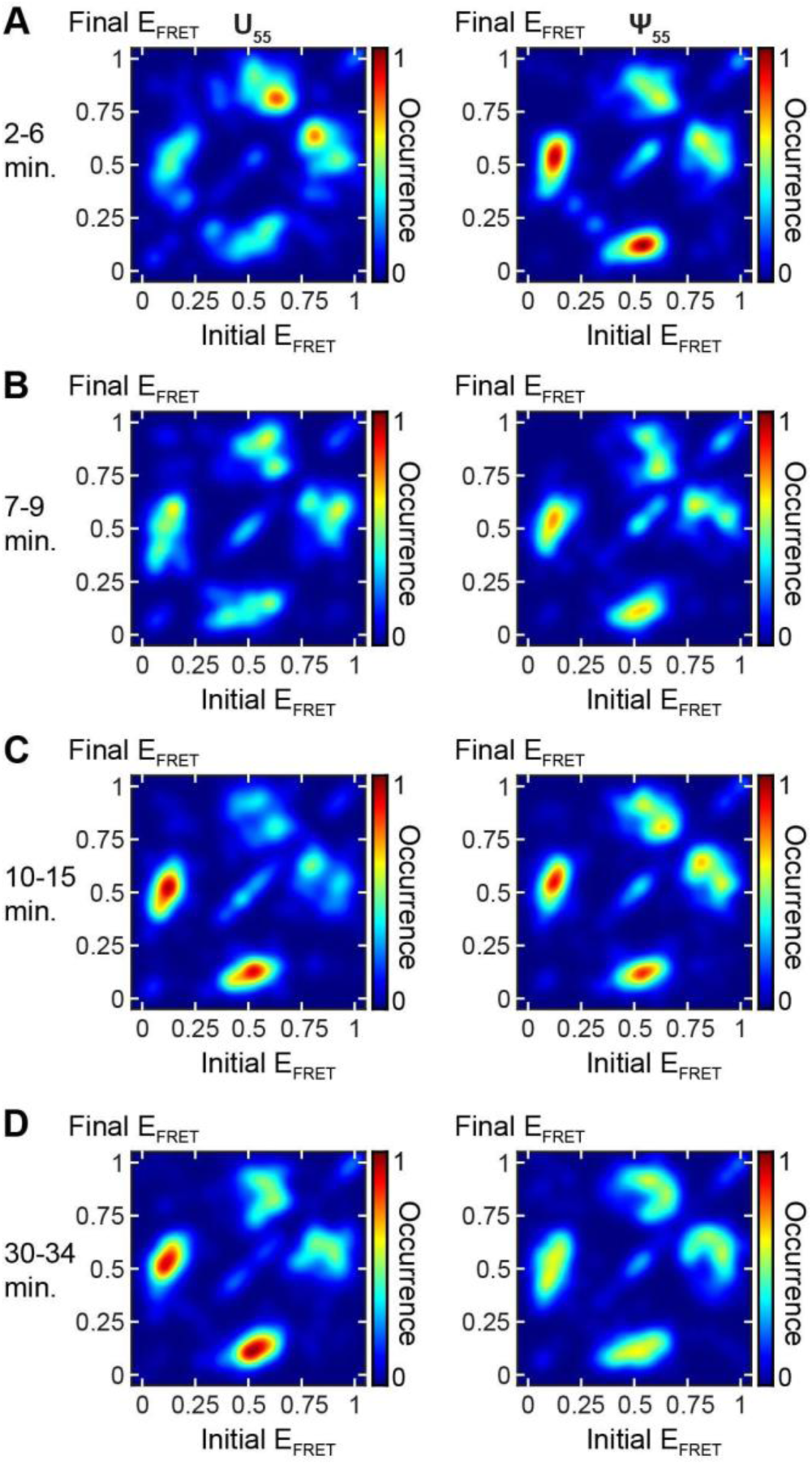
TODPs for U_55_ and Ψ_55_ tRNA eMet in the presence of 200 nM Pus4. TODPs for U_55_ (left column) and Ψ_55_ (right column) tRNAs compiled from traces recorded (*A*) 2–6 minutes after injection of protein; (*B*) 7–9 minutes; (*C*) 10–15 minutes; (*D*) 30–34 minutes.

As a control, we binned smFRET data recorded on U_55_ tRNA in the absence of Pus4 into the same time windows (**Fig. 7C, Supplemental Fig. S12**). We observed small changes in the relative amplitudes of the histogram features, particularly LF and HF. These changes were likely due to the finite number of molecules analyzed for each time window, and the gradual activation and depletion of the oxygen scavenging system used to prolong fluorophore activity. Importantly, however, U_55_ tRNA alone did not exhibit the curve broadening, shifts in E_FRET_, or increase in overall intensity (**Supplemental Table S2**) that had characterized its time evolution in the presence of Pus4.

To help isolate the influence of Pus4 binding and remodeling of tRNA from its pseudouridylation activity, we performed smFRET experiments with the pre-modified Ψ_55_ tRNA eMet in the presence of Pus4. While MF microstates were observed in Ψ_55_ tRNA eMet at early times (**Supplemental Fig. S11B**), as we also observed for U_55_ tRNA eMet, Ψ_55_ tRNA histograms and TODPs exhibited less dramatic changes. Specifically, in the 2–6 minute window following injection of Pus4, Ψ_55_ tRNA exhibited a smaller shift of the MF state toward higher E_FRET_ than U_55_, and an increase in the population of the HF state (**Fig. 6A**). Ψ_55_ tRNA retained the same shape in its histogram throughout the remainder of the time course, exhibiting only minor fluctuations of the relative populations of its states (**Fig. 6**, **Fig. 7B)**. These fluctuations were of comparable size to those observed in tRNA alone (**Fig. 7C, Supplemental Fig. S12)**, so we conclude that the impacts of Pus4 on the conformational ensemble of Ψ_55_ tRNA are largely complete after 2–6 minutes of interaction. This stands in contrast to U_55_ tRNA, which continues to evolve through at least 15 minutes after injection of Pus4 despite the protein binding at a similar rate (as indicated by BLI and PIFE) and Pus4 presumably pseudouridylating most U_55_ tRNA molecules within 2 minutes (**Fig. 2**). By 10–15 minutes of incubation with Pus4, U_55_ tRNA exhibited only a small skew toward lower E_FRET_ relative to Ψ_55_, and their histograms were nearly superimposable by 30–34 minutes (**Fig. 6C,D**). We hypothesize that Ψ_55_ tRNA thus bypassed some of the conformational changes that U_55_ tRNA transitioned through during its catalysis and subsequent remodeling by Pus4.

## Discussion

In the decades since the three-dimensional structure of tRNA was elucidated (Kim et al. 1974), it has been presented as an archetype of structured non-coding RNA. Yet every tRNA undergoes many processing steps during biogenesis—including trimming, tailing and the addition of multiple chemical modifications—necessitating many conformational changes. In this study we combined biophysical and biochemical approaches to document the dynamics of conformational changes that two yeast tRNAs can undergo over time. We then measured how this chronology of tRNA structural dynamics changed upon binding by a ubiquitous pseudouridine synthase that modifies many tRNAs in nature. These data demonstrated how Pus4 impacts tRNA conformation through its binding and catalysis, leading to a more well-defined ensemble of conformations when the RNA-protein complex reached equilibrium. This outcome was reached after a period of minutes if catalysis was bypassed (i.e. Pus4 incubated with a pre-modified tRNA), rather than *tens* of minutes if catalysis occurred.

We find that tRNA eMet and tRNA Thr each sample complex free energy landscapes, with at least three conformational states resolved by smFRET. Both exhibit a hierarchical folding landscape, with their MF states as intermediates, though tRNA Thr exhibits faster dynamics. Furthermore, tRNA eMet maintains a dynamic conformational landscape of LF, MF and HF states throughout its interactions with Pus4 and independently of its modification status, with additional MF states populated upon engagement with Pus4. In the absence of protein, the E_FRET_ values of these features align reasonably closely with FRET states previously observed in human mt-tRNA^Lys^ under equivalent ionic conditions of 50 mM buffer, 100 mM monovalent cation and no divalent cation (Dammertz et al., 2011). However, we consistently observe a larger HF population in the yeast tRNA eMet than previously observed in mt-tRNA^Lys^, which we propose can be reconciled by predicted free energies. The cloverleaf secondary structure of tRNA eMet has a predicted ΔG_folding_ = −18.9 kcal/mol (details of free energy calculations in **Methods**), significantly more stable than −16.1 kcal/mol for mt-tRNA^Lys^ studied in Voigts-Hoffman *et al*., 2007 and subsequent studies (Dammertz et al. 2011; Kobitski et al. 2011, 2008) (**Supplemental Fig. S13A,E**).

mt-tRNA^Lys^ also substantially populated an MF state with E_FRET_ ∼0.4 that interconverted with C on a time-scale of hundreds of milliseconds (Voigts-Hoffmann et al. 2007). This state was assigned to an extended hairpin secondary structure (E) that had previously been identified in this tRNA by biochemical methods (Helm et al. 1998). Interconversion between C- and E-type structures was also observed using biochemical methods in a different tRNA, mt-tRNA^Leu^ (Bhaskaran et al. 2014). Consistent with these experimental observations, the E structure of mt-tRNA^Lys^ is predicted to have a free energy of −15.3 kcal/mol, within 1 kcal/mol to that of the C structure (**Supplemental Fig. S13E,F**). To place our results into this context, we constructed a plausible E-like secondary structure of yeast tRNA eMet by identifying potential base pairs that could be formed between nucleotides in the T- and D-loops and adjacent stems (**Supplemental Fig. 13B**). Unlike mt-tRNA^Lys^, however, at ΔG = −12.9 kcal/mol, this E-like structure of tRNA eMet is predicted to be 6 kcal/mol less stable than its C structure (**Supplemental Fig. 13A,B**). Likewise, tRNA Thr is predicted to have a C structure that is 7 kcal/mol more stable than a plausible E structure (−24.4 and −17.4 kcal/mol, respectively) (**Supplemental Fig. 13C,D**). In light of the substantial free energy preference for the C secondary structures of tRNA eMet and tRNA Thr, we consider it unlikely that the dominant MF states that we observe both with and without Pus4 arise from E-like secondary structures. Instead, the MF states observed in tRNA alone and the variations thereof that are populated in tRNA eMet in the presence of Pus4 may be explained by 1) relatively minor changes in secondary structure such as fraying/unpairing of the acceptor stem that contains the donor fluorophore, and/or 2) changes in tertiary structure such as breaking of the T-loop-D-loop interaction. These structures have been observed in *E. coli* tRNAs *in vivo* under stress conditions (Yamagami et al. 2022). Another recent study that used single particle cryo-EM and melting temperature assays found that tRNA folding was strongly influenced by the tertiary structure stabilizing interaction between the D- and T-arms, and that this interaction itself was generally promoted by Ψ_55_ (Biela et al. 2025). Prior steady-state and kinetic measurements also identified multiple interconvertible tertiary structures under conditions generally favoring the C secondary structure (Cole and Crothers 1972). Future studies of additional tRNAs and models with alternate placement of FRET pairs/fluorophores may improve delineation of the structures that give rise to the observed FRET states.

Our work herein began with the goal to characterize the conformational dynamics of tRNA as influenced by pseudouridylation and interaction with a pseudouridine synthase. Guided by our results, we propose that each instance of Pus4 engaging with tRNA follows a three-to-four-step process: 1) Initial binding of Pus4 occurs, 2) Pus4 “tests” whether Ψ_55_ is already present, 3) If Ψ_55_ is absent, Pus4 may convert U_55_ to Ψ_55_, 4) Pus4 remodels tRNA into a new distribution of structures. Pus4 effectively discriminates U from Ψ, remodeling pre-modified tRNA toward the final structural ensemble within minutes. Upon initial engagement with unmodified or pre-modified tRNA, Pus4 induces an additional population of intermediate (MF) states. Such structural states may play a role in discrimination between substrate and product.

Overall our results demonstrate two important layers of influence on tRNA conformational dynamics: 1) chemical catalysis, and 2) remodeling resulting from extended and recurrent interactions with modification enzymes. Why Pus4 would promote such malleability in tRNA conformational dynamics on already pseudouridylated tRNAs remains an important question. Since Pus4 catalysis can strongly influence the addition of subsequent modifications *in vivo* (Shaw et al. 2024), and some are added over many hours after pseudouridylation (Barraud et al. 2019), we propose that part of the role of Pus4 is to make tRNA structure amenable to binding and modification by subsequent modification enzymes that are critical for tRNA function.

## Materials and Methods

### Synthetic tRNA sequence design

Fluorophore placement was modeled after that used in Voigts-Hoffmann *et al*., 2007. The yeast tRNA eMet and tRNA Thr were chosen for these reasons: 1) both contain uridines at positions 4 and 41, which were the positions labeled with fluorophores and match those labeled in the prior work (Voigts-Hoffmann et al. 2007), 2) both have a minimum free energy structure, as predicted by RNAFold (Chan and Lowe 2009, 2016), that closely resembles the tRNA cloverleaf secondary structure, theoretically reducing formation of non-canonical, off-pathway structures *in vitro*, and 3) prior evidence demonstrates that both tRNAs are pseudouridylated by Pus4 efficiently *in vivo* (Shaw et al. 2024), while our work herein demonstrated their efficient pseudouridylation by Pus4 *in vitro*. The tRNAs were designed to contain Cy3 attached to the carbon-5 position of U_4_ and Cy5 attached to the carbon-5 position of U_41_, as well as biotinylation of the 5′-phosphate to enable attachment of the molecules to the microscope slide, or probe, for smFRET and BLI, respectively.

### Ligation of synthetic tRNA sequences

Full-length tRNA constructs were made by ligation of two RNA oligonucleotides (Dharmacon, HPLC-purified by the manufacturer) using a DNA splint (Integrated DNA Technologies, PAGE-purified by the manufacturer) (**Supplemental Table S3**). Oligonucleotide concentrations were quantified using A260 measurements on a Nanodrop spectrophotometer and extinction coefficients provided by the manufacturer. Ligation was performed using 2 nmol of each of the 5’ and 3’ RNA segments and splint in a combined volume of 60 µl to which 7.5 μL hybridization buffer (150 mM HEPES pH 7.5 and 500 mM KOAc) was added. This solution was heated for 2 minutes at 90°C and allowed to cool at room temperature for 30 seconds before 10X T4 RNA ligase 2 buffer (NEB) was added to a final concentration of 1X. The mixture was then cooled at room temperature for an additional 10 minutes, at which time additional ligase buffer, ATP, water, and T4 RNA ligase 2 (MC Labs) were added to achieve a reaction volume of 100 μL containing 0.4 mM ATP, 1X RNA ligase 2 buffer, and 100 units T4 RNA ligase 2. The ligation mixture was incubated at room temperature for 3 hours. Following this, 16 units DNase I (NEB) was added along with 10X DNase I buffer to bring the final concentration of that buffer to 1X. This mixture was incubated at 37°C for 15 minutes to digest the DNA splint.

A 2X FALB denaturing loading buffer consisting of 90% Formamide (TCI) in 1X TBE was added to a final concentration of 1X and the ligation mixture was heated for 2 minutes at 90°C to inactivate the enzyme and unfold the RNA. The resulting RNA mixture was run on a 12% polyacrylamide, 8 M urea denaturing gel and imaged using an Amersham Typhoon (GE). The gel was then stained with Sybr gold in 1X TBE for 15 minutes and the 76 nt band corresponding to the full length tRNA was excised while visualizing the gel on a Dark Reader transilluminator (Clare Chemical Research). The RNA was extracted using an electro-elution system (BioRad) run at 10 mA per fritted tube for 15 minutes in 0.5X TBE. After elution, the polarity was reversed for 1 minute to release the RNA from the membrane and the product was recovered from the apparatus. 1/10 volume of 3 M sodium acetate pH 5.5 and 2.5 volumes of −20°C 100% EtOH were added and the solution was mixed and stored at −20°C overnight. Tubes were centrifuged at 4°C, 10,000 rcf for 40 minutes. The supernatant was removed, 500 μL of −20°C 70% EtOH was added, the tube was vortexed for 30 sec, and then centrifuged at 4°C, for 10 minutes at the maximum RCF rating of the tube. The supernatant was again removed and pellets were dried in a vacuum centrifuge at room temperature for 15 minutes until dry. The pellets were dissolved in 50 μL DEPC-treated H2O and quantified using a NanoDrop.

### Protein expression and purification

A pGEX-5X-3-ScPus4 backbone plasmid encoding GST-Pus4 with a Factor Xa cleavage site between the GST tag and Pus4 protein sequence (PDG322) was used (**Supplemental Table S1, S3**). The GST-tagged Pus4 vector was transformed into *E. coli* BL21(DE3) competent cells (Thermo Fisher Scientific). A single transformant of each was inoculated to make a 50 mL overnight starter culture, which was then diluted into a 4 L culture grown until reaching OD_600_=0.80. Cultures were grown at 37°C in TB medium containing 100 µg/mL ampicillin and shaken at 225 rpm. Upon reaching the desired OD_600_, the 4 L culture was shifted to 16°C for 30 minutes. Protein expression was induced by adding IPTG (Gold Bio) to a final concentration of 1 mM, and cells were grown overnight for ∼16 hours at 16°C with continuous shaking at 225 rpm. Cells were harvested by centrifugation, and the cell pellets were resuspended in 30 mL of Buffer A (50 mM Tris-HCl pH 7.5, 400 mM NaCl, 10 mM MgCl_2_, 10% Glycerol, 0.5 mM EDTA, 5 mM BME, 1 tablet of Pierce EDTA-free Protease Inhibitor (Thermo Fisher Scientific)), divided into two 50 mL conical tubes, and then flash-frozen in liquid nitrogen before storage at −80°C. Each tube contained enough material for a single protein purification. All subsequent procedures were performed at 4°C unless otherwise stated.

Conical tubes were thawed on ice. The following reagents were added to achieve their final concentrations, respectively: lysozyme (Sigma) to 1 mg/mL, DNase (GoldBio) to 10 µg/mL, MgCl_2_ (Fisher) to 0.5 mM, and RNase A (Fisher) to 30 µg/mL. This cell suspension was incubated for 30 minutes. Cells were lysed via sonication using a sonicator (Branson Sonifier 450) with 30-seconds-on/30-seconds-off cycles (50% duty cycle, power level of 4), repeated three times. Insoluble material was removed by centrifugation for 30 minutes at 30,000 x g in a JA-20 rotor. After clarification, the remaining soluble material was incubated with 2 mL of a 50% slurry of Glutathione-Sepharose beads (Cytiva) that had been pre-equilibrated with Buffer A for 1 hour with end-over-end rotation at room temperature. After incubation, the protein-bound beads were isolated using centrifugation (5 minutes at 500 x g) and then washed with 20 mL of Buffer A. This wash step was repeated for a total of five times. Protein was then eluted from the beads with 1 mL of freshly prepared Elution buffer (50 mM Tris-HCl pH 7.6, 10 mM reduced glutathione) after incubating for 5 minutes at room temperature with end-over-end rotation. This elution step was repeated five times and the elutions were pooled together (∼5 mL).

The eluate was concentrated to ∼1 mL final volume using a 30 kDa Vivaspin concentrator (GE Healthcare). Using the same concentrator, the sample underwent exchange into Buffer B (50 mM Tris-HCl pH 7.5, 100 mM KCl) by adding 10 mL of Buffer B, and performing centrifugation to concentrate to ∼1 mL final volume. This was repeated two more times. These protein samples were aliquoted into Eppendorf tubes and flash frozen in liquid nitrogen before storage at −80°C. The yield was quantified using A_280_ measurements on a Nanodrop spectrophotometer. Protein purity was determined using SDS-PAGE with samples collected across steps of the protein purification process (**Supplemental Fig. S1A**).

### Testing purified protein for RNase contamination

Approximately 1 µg of tRNA isolated from yeast was incubated with 0.4 µM of Pus4 in a 10 µL total volume for one hour at room temperature. Control samples of tRNA without protein and tRNA plus protein without incubation were also prepared. Samples were combined with 10 µL of 2X RNA loading dye (NEB), heated at 70°C for 8 minutes on a heat block, and then loaded onto a fresh pre-run PAGE gel, along with RNA ladder, using an Apogee V16 gel system. After running the gel at 60 mA for about 1.5 hours, the gel was stained with SYBR Gold for 10 minutes at room temperature, and imaged using an Amersham Typhoon (GE) to assess tRNA degradation (**Supplemental Fig. S1B**).

### Yeast growth conditions

*pus4*Δ (YDG127) cells were streaked out onto YPD agar media plates from a frozen −80°C stock and grown at 30°C for 2 days (**Supplemental Table S1**). Colonies were inoculated into 5 mL YPD media in culture tubes and grown at 30°C in a rolling drum overnight. In the morning, 400 μL of each culture was transferred to culture tubes with 5.2 mL liquid YPD and grown at 30°C in a rolling drum to an OD_600_ of ∼0.8. Cultures were transferred to 15 mL conical tubes and centrifuged at 4°C, 3000 x g for 2 minutes. Media was decanted, and the cell pellets were resuspended in 1 mL cold 1X PBS and transferred to microcentrifuge tubes. Tubes were centrifuged at 4°C, 9500 rpm for 1 minute to pellet. Supernatant was decanted and cell pellets flash frozen by placing microcentrifuge tubes in liquid nitrogen. Cell pellets were stored at −80°C until used for total RNA isolation.

### tRNA isolation from yeast

*pus4*Δ (YDG127) yeast pellets were thawed on ice (**Supplemental Table S1**). For cell lysis and RNA preservation, pellets were resuspended in 400 μL TES buffer (10 mM Tris, 10 mM EDTA, 20% SDS). 400 μL acidic phenol was added to each tube. Tubes were vortexed for 10 seconds and incubated in a 65°C water bath for one hour, with 10 seconds of vortexing every 15 minutes. Tubes were put on ice for 5 minutes then centrifuged at 4°C, 13,000 rpm for 10 minutes. The top aqueous layer of each sample was transferred to a new tube. 400 μL of chloroform was added to each of the new tubes. Tubes were vortexed for 10 seconds then centrifuged at 4°C, 13,000 rpm for 10 minutes. The top aqueous layer of each sample was transferred to a new tube and the addition of chloroform, vortexing, and centrifuging repeated. The top aqueous layer of each sample was transferred to a new tube. To precipitate RNA, 1/10 volume of 3 M sodium acetate pH 5.5 and 2.5 volumes of ice cold 100% EtOH were added to each sample and mixed. Tubes were stored at −80°C for one hour. Tubes were centrifuged at 4°C, 13,000 rpm for 10 minutes. The supernatant was removed and 500 μL cold 70% EtOH added to each tube to wash pellets of salt. Tubes were centrifuged at 4°C, 13,000 rpm for 10 minutes. Ethanol was removed and tubes left open for 5–10 minutes to allow residual ethanol to evaporate. Dried pellets were resuspended in 25 μL water, and their concentrations were quantified using a Nanodrop.

An 8% urea acrylamide gel was made using the National Diagnostics UreaGel 19:1 Denaturing Gel System. The gel was pre-run in 1X TBE running buffer at 45 mA for 30 minutes. Total RNA samples were aliquoted to 10 μL volumes and 10 μL 2X RNA loading dye (NEB) added to each sample. 3 μL Low Range ssRNA Ladder (NEB) was mixed with 7 μL water and 10 μL 2X RNA loading dye. Samples were incubated at 70°C for 8 minutes. Samples were loaded on the gel and empty wells loaded with 10 μL water + 10 μL 2X RNA loading dye. The gel was run at 20 mA for 15 minutes and then at 60 mA for ∼1 hour. The gel was stained with SYBR gold (Invitrogen) in 1X TBE running buffer for 10 minutes. The gel was imaged with an Amersham Typhoon (GE). Using UV shadowing, tRNA bands were cut from the gel. Gel fragments were placed in tubes with 450 μL 0.3 M NaCl. Tubes were put on a tube inverter overnight at 4°C. Liquid was transferred to new tubes and 1.05 mL 100% EtOH was added to each sample and mixed. Samples were stored at −80°C for one hour. Samples were centrifuged at 4°C, 13,000 rpm for 35 minutes. The supernatant was removed and tubes left open for 5–10 minutes to allow residual ethanol to evaporate. Dried pellets were resuspended in 10–20 μL water and stored at −80°C.

### *in vitro* modification

To perform the *in vitro* modification, 1 µM Pus4 protein was mixed with 300 nM total tRNA harvested from *pus4*Δ cells and incubated at 30°C for 2 minutes in a 200 μL reaction volume in Buffer B (50 mM Tris-HCl pH 7.5, 100 mM KCl). Buffer B supplemented with 4 mM MgCl_2_ was also tested to observe whether magnesium affected catalysis. Since it did not have a strong effect (**Fig. 2**), MgCl_2_ was excluded from the buffer used for all smFRET and BLI assays. These reactions were halted with the addition of 200 µL of saturated phenol (pH 6.6). Samples were mixed and then stored at −80°C.

To purify *in vitro* modified tRNA, samples were thawed on ice, then vortexed for 10 seconds and centrifuged at 4°C, 13,000 rpm for 10 minutes. tRNA was re-purified as done in the tRNA isolation (described above), beginning with the addition of 400 μL chloroform. Samples were resuspended in 10 μL water and concentrations quantified using a Nanodrop. Samples were stored at −80°C.

### Direct sequencing of *in vitro* modified tRNA

To perform library preparation for sequencing using Oxford Nanopore Technology (ONT), samples were first thawed on ice. Samples were incubated at 95°C for 2 minutes, room temperature for 2 minutes, then on ice for 2 minutes. In a new tube, 7.25 μL tRNA sample was mixed with 2 μL 10X RNA ligase 2 buffer (NEB), 0.8 μL each of: 10 μM ACCA, 10 μM GCCA, 10 μM UCCA, 5 μM CCCA, and 5 μM CCA specific adapters (Shaw et al. 2024), 2 μL 50% PEG, 2 μL 250 mM MgCl_2_, 1.25 μL 100 mM DTT, 2μL 20 mM ATP, and 1 μL RNA ligase 2 (NEB), for a total volume of 20 μL. The reaction was incubated at room temperature for 45 minutes. 20 μL water was added to the reaction. tRNA was purified with 1.8X volume RNAClean XP Beads (Beckman Coulter). Samples were washed twice with 500 μL 80% EtOH, eluted with 23 μL water. Samples were mixed with 8 μL 5X quick ligase buffer (NEB), 6 μL RMX adapter (ONT), and 3 μL T4 DNA ligase (2,000,000 U/mL; NEB) for a total volume of 40 μL. The reaction was incubated at room temperature for 30 minutes. tRNA was purified with 1.5X volume RNAClean XP Beads. Samples were washed twice with 150 μL WSB (ONT), then eluted with 15 μL ELB (ONT). Water was added to bring the total volume to 37.5 μL. Samples were sequenced with MinION devices following the ONT SQK-RNA002 protocol.

Pseudouridylation of U_55_ in tRNA leads to a base miscall, due to an alteration in the ionic current when a pseudouridine transitions through the nanopore, as compared to the different disruption in ionic current caused by the unmodified uridine base. This causes a “mismatch” between the sequenced base and the unmodified reference base. The “reference match probability” refers to the probability that a given base will match the reference base. Therefore, a low reference match probability at position 55 is indicative of a pseudouridylated base. For more details see Shaw and Thomas et al. 2024.

Sequencing results were basecalled using Guppy v3.0.3, aligned with BWA-MEM (-W 13 –k 6 – xont2d), and quality controlled with SAMtools (Q1). marginAlign was used to create alignment and error models, and marginCaller was used to calculate mismatch probabilities. Python was used to calculate the reference match probability—equal to a value of 1.0 minus the mismatch probability value—and generate heatmaps. DRS data files are listed in **Supplemental Table S4**.

### Preparation of sucrose gradients

A 6X Buffer B solution was prepared, then mixed with a 60% sucrose solution dissolved in water at the appropriate ratios to achieve five sucrose solutions of 10%, 15%, 20%, 25%, and 30%—each achieving a final concentration of 1X Buffer B. Once prepared, 980 µl of the 30% sucrose solution was pipetted into Ultra-Clear centrifuge tubes (½ inch x 2 inch, Beckman Coulter catalog #344057), then allowed to freeze at −80°C for 20 minutes. The 25% solution was layered on top, allowed to freeze, and this was repeated for the remaining solutions. Any samples to be compared were run with the same batch of sucrose gradients to control for batch variation. Sucrose gradients were prepared ahead of time and stored at −80°C.

### Sucrose gradient ultracentrifugation

Gradients were thawed overnight at 4°C prior to use the next day. A 100 µL solution with 10 µM purified GST-Pus4 with and without 10 µM tRNA derived from YDG1 (**Supplemental Table S1**) was allowed to sit at room temperature for 30 minutes prior to loading onto a gradient. Gradients were balanced, and then spun in an ultracentrifuge (Beckman Coulter Optima XPN-100) in a SW 55 Ti swinging bucket rotor (Beckman Coulter) for 4 hours at 55,000 rpm. Following ultracentrifugation, gradients were fractionated by hand, serially removing 200 µl from the top of the gradient and aliquoting into 24 tubes. (Note that the final fraction had a volume in the range of 200–400 µl, due to subtle differences in the total gradient volume and aspiration efficiencies of the dense sucrose mixture after spinning). Fractions were flash frozen in liquid nitrogen prior to storage at −80°C.

Five nanomoles of each gel filtration chromatography-purified standard (beta-amylase and carbonic anhydrase, Millipore Sigma) were combined and run together in a separate gradient.

### SDS-PAGE of sucrose gradients

Sucrose gradient fractions were mixed 1:1 with 2X Laemmli buffer (Bio-Rad) containing 5% v/v 2-mercaptoethanol, vortexed for several seconds, boiled on a heat block at 95°C for 10 minutes, then vortexed again and briefly spun down to condense liquid. Equal volumes from each fraction were loaded across three 4–15% PAGE gels (Bio-Rad). After electrophoresis in 1X SDS running buffer, gels were stained with GelCode Blue stain reagent (ThermoFisher Scientific) to visualize protein abundance across the fractions.

### Biolayer Interferometry (BLI)

BLI was performed using the biotinylated and fluorophore-labeled RNA substrates used for smFRET as well as versions of tRNA eMet lacking fluorophores. After the ligation steps described above, the RNA was diluted in Buffer B to a concentration of 60 nM. The solution was heated to 90°C on a heat block for 2 minutes, then allowed to cool for 10 minutes at room temperature before placing it on ice. BLI measurements were performed at 30 °C on a GatorPlus Biolayer Interferometer (GatorBio, Palo Alto, CA) utilizing proprietary GatorOne software for data collection. GatorBio streptavidin-coated glass probes were presoaked in Buffer B for at least 20 minutes before collecting baseline signal for 300 seconds. The probes were immersed in 60 nM biotinylated tRNA until an adequate shift (∼1.2 nm) in signal was observed (∼60 seconds). The probes were then submerged in 10 nM biocytin (Cayman Chemical) in Buffer B to quench the remaining streptavidin sites. This was followed by a second quenching in Buffer B supplemented with 0.2% Tween-20, which further aided in reducing non-specific protein binding. For this reason, Buffer B supplemented with 0.02% Tween-20 (“Buffer C”) was used for all following steps. Loaded probes were then incubated in Buffer C until baseline was achieved (∼300 seconds). To measure association of the protein to the tRNA, the loaded probes were then incubated in 200 nM of GST-Pus4 in Buffer C for ∼2500 seconds or more. To measure dissociation, the probes were then submerged in Buffer C for the same amount of time as the association step. Reference probes were included in some runs to assess non-specific protein binding and RNA dissociation from the probe.

### Analysis of BLI data

To estimate the kinetics of Pus4 binding to tRNA, we attempted to isolate the initial rate of association before pseudouridylation (in the case of tRNA that was not premodified) or refolding occurred. In isolation, this early phase of binding represents a reversible second order reaction (RNA+Pus4 ⇋ RNA·Pus4). We approximated its kinetics as pseudo-first order in Pus4, as the protein was present in bulk solution while the quantity of tRNA was limited by the number of binding sites on the probe. This is expected to yield an exponential shape to the association curve (**Eq. 1**):

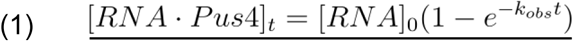

where the observed rate of the reaction is:

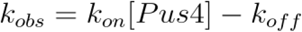

Considering that at each timepoint *t*, [RNA]_0_ = [RNA·Pus4]_t_ + [RNA]_t_, and that at the final timepoint [RNA]_final_ = [RNA]_0_ - [RNA·Pus4]_final_, **Eq. 1** becomes:

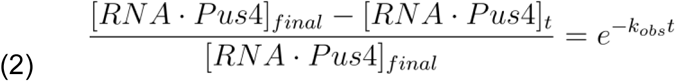

To estimate *k_on_* and *k_off_*, the association phases of the BLI curves were converted to fractional saturation (FS) using **Eq. 3**, where *s(t)* is the shift at time *t* and *s_final_* is the shift at the final timepoint of the association phase:

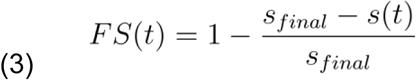

Two methods were then used to fit the fractional saturation curve: 1) The early segment of the association phase over which ∼75% of the overall shift occurred was fitted with a single exponential (**Eq. 4**), or 2) a longer period of the association phase was fitted with the sum of 2 exponentials (**Eq. 5**). The first 1000 s were used in the bi-exponential fits for tRNA without fluorophores due to the prominent downturn in the curves at 800 nM, while the entire association phase was fitted in all other cases. In **Eq. 5**, the faster of the two rates was taken to be *k_obs_*.

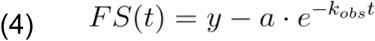

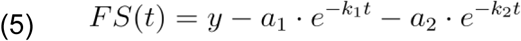

These fits were performed for BLI curves obtained at 50 nM, 200 nM and 800 nM Pus4. *k_obs_* was then plotted against [Pus4] and the slope of a linear fit yielded an estimate for *k_on_*. In principle, the y-intercept of the fit yields *k_off_*. However, determining it accurately would require measurements at lower [Pus4], conditions that reduce both the accuracy of the pseudo-first order kinetics approximation and the separation of the timescales on which Pus4 accumulates on the probe and on which it modifies and/or remodels the tRNA it has already bound to. Because *k_obs_* was less than 0.01 s^-1^ at 50 nM Pus4 (the lowest concentration tested) for all RNAs and fitting approaches, we can reasonably assume that *k_off_* is less than that value. Given the values obtained for *k_on_* (**Supplemental Fig. S5**), this constrains the dissociation constant *K_D_ = k_off_/k_on_* to a value less than ∼130 nM.

### Electrophoretic Mobility Shift Assay

The RNA was annealed as described above for BLI at a concentration of 300 nM. The RNA was then further diluted to 30 nM in Buffer B and incubated with varying concentrations (0 nM, 15 nM, 30 nM, 60 nM, 120 nM, 200 nM, 400 nM, 600 nM, 800 nM) of Pus4 in a total volume of 10 μL for 30 minutes at room temperature. Each 10 μL solution was mixed with 10 μL of loading buffer (2X TBE, 40% glycerol). The mixtures were run on an 8% Polyacrylamide native/non-denaturing gel at 300 V for 2.5 hours at 4°C. The gel was prepared from 40% 19:1 Acrylamide/Bisacrylamide (Bio-Rad), TEMED (Thermo Fisher Scientific), and Ammonium Persulfate (Thermo Fisher Scientific) and pre-run in 1X TBE running buffer at 300 V for 90 minutes. The gel was imaged using an Amersham Typhoon (GE).

### Single molecule FRET (smFRET)

RNA was annealed as described above at a concentration of 10 nM, then diluted to 10 pM in Buffer B supplemented with 5 mM D-glucose (Sigma Aldrich), 165 U/mL glucose oxidase from *Aspergillus niger* (Sigma Aldrich), 2150 U/mL catalase from *Corynebacterium glutamicum* (Sigma Aldrich) and 5 mM Trolox (Sigma Aldrich) as an oxygen scavenging system (OSS). This OSS was used to visualize the RNA during immobilization due to its fast acting capability. This mixture was flowed onto a slide that had been passivated with DDS (Sigma Aldrich) as previously described (Hua et al. 2014), incubated with 0.2 mg/mL BSA (Thermo Fisher Scientific) for 5 minutes, 0.2% v/v Tween-20 for 10 minutes, and 0.2 mg/mL streptavidin (Thermo Fisher Scientific) for 5 minutes. Imaging was performed with Buffer C containing 5 mM 3,4-dihydroxybenzoic acid (Sigma Aldrich), 5 mM Trolox (Sigma Aldrich), and 50 nM Protocatechuate 3,4-dioxygenase from *Pseudomonas sp.* (Sigma Aldrich or MP Biomedicals) as an OSS and, for experiments with protein, 200 nM Pus4. This imaging buffer was incubated in a tube with minimal headspace for 10 minutes to allow the OSS to begin depleting oxygen before injection into the sample chamber. Data were collected at room temperature on a home-built prism-type TIRF microscope (Leica DMi8 inverted microscope, 63X oil immersion objective). Cy3 and Cy5 were excited by a 532 nm laser (Coherent OBIS 532 LS) at 140 mW and a 637 nm laser (Coherent OBIS 637 LX) at 60 mW, respectively. The slide was illuminated at 532 nm continuously and at 637 nm at the beginning and end of each movie to confirm that Cy5 was active. 2-minute-long movies were recorded on an Andor iXon 888 EMCCD with an exposure time of 100 ms. An EM gain setting of 300 was used for movies recorded in the presence of Pus4. tRNA-only movies used for intensity comparison (**Supplemental Table S2**) were recorded at the same EM gain, whereas tRNA-only movies used for FRET analysis (**Fig. 5**, **Supplemental Fig. S12**) were recorded at an EM gain of 450 due to lower fluorophore brightness in the absence of protein. Movies were collected back to back for the first 15 minutes following injection of imaging buffer, and additional movies were collected 30–34 minutes after injection. Unless otherwise noted, data reported for tRNA in the absence of Pus4 was recorded for 10–15 minutes following injection of imaging buffer.

Coordinates for each molecule were extracted from raw video files using FIJI and fluorophore intensity traces were extracted using custom MATLAB code. Traces that exhibited single-step photobleaching of Cy3 and Cy5 and anticorrelation between donor and acceptor signals when transitions occurred were selected manually and analyzed using custom MATLAB codes. FRET efficiency was calculated from the measured donor and acceptor emission intensities (I_D_ and I_A_) using **Eq. 6**:

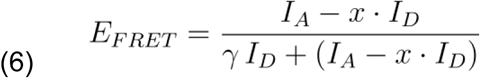

Variable x corrects the acceptor signal for bleedthrough of donor emission into the acceptor channel and was determined to be 0.129 through analysis of tRNA molecules that lacked an active Cy5. γ was determined through analysis of traces in which Cy5 photobleached before Cy3, as previously described (McCann et al. 2010), and was found to be 2.42 in the absence of protein and 3.01 in the presence of protein (**Supplemental Fig. S8C,D**). Noise in the corrected donor and acceptor traces was suppressed using a bilateral filter with a spatial spread of 2 frames and a signal value spread of approximately 1/4 of the mean signal intensity across all traces in a dataset for Cy3 and approximately 1/5 of the mean signal intensity for Cy5. This resulted in signal value spreads of 900–1000 for Cy3 and 800–1100 for Cy5 in the absence of protein, and spreads of 2200–3100 for Cy3 and 1900–2500 for Cy5 in the presence of protein, which preserved rapid excursions in the resulting E_FRET_ traces.

FRET histograms were compiled from the first 50 frames of ∼200 traces and fitted with a sum of three or four Gaussian functions in Mathematica (Wolfram Alpha). Hidden Markov Modeling was performed in QuB (Quantifying unknown Biophysics) Online (Nicolai and Sachs 2013; Milescu et al. 2000; Qin 2004) and TODPs were generated from the resulting idealized traces using custom MATLAB codes. Correlation analysis was performed in Mathematica by computing the cross-correlation C(τ) between Cy3 and Cy5 intensities using **Eq. 7** (Kuzmenkina et al. 2005).

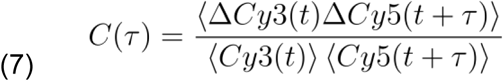

where Δ*X*(*t*) = *X*(*t*) − 〈*X*〉 and 〈*X*〉 is the average value of X across all frames t within the trace. For autocorrelation analysis, Cy3 and Cy5 intensities in the above equation were both replaced by the E_FRET_ value. Traces used for correlation analysis were corrected for bleedthrough and γ factor but not subjected to the bilateral filter. Autocorrelation and cross-correlation curves were fitted simultaneously with a two-exponential model (**Eq. 8 and 9**, respectively) with the time constants t_1_ and t_2_ fixed across the two curves.

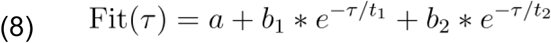

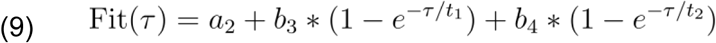

HMM-derived rate constants for transitions between FRET states were determined by extracting the durations of dwells in each state and fitting their cumulative distribution functions with **Eq. 9**, where k_i_=1/t_i_.

### Bulk emission spectroscopy

Emission spectra were recorded on an Edinburgh Instruments FS5 spectrofluorometer using a 4 mm square cuvette. RNA was annealed as described above at a concentration of 150 nM in Buffer B and used without further dilution. Cy3 was excited at 532 nm with a bandwidth of 2 nm and emission was scanned from 500 to 800 nm in 1 nm steps with a bandwidth of 2 nm and a 1 s dwell time.

### Free energy calculations

The free energies of different secondary structures were predicted using the RNAfold web server (Gruber et al. 2008). The sequence used for mt-tRNA^Lys^ was the one utilized in Voigts-Hoffman *et al*., 2007 with U instead of dT at its fluorophore labeling sites, and the sequence used for tRNA eMet was the one utilized in this work. Mild base-pairing constraints were applied as needed to achieve the C and E secondary structures of mt-tRNA^Lys^ as depicted in Fig. 1 of Voigts-Hoffman *et al*., 2007, the canonical tRNA eMet and tRNA Thr structures as depicted in **Fig. 1A** of this work, and plausible E-like structures of tRNA eMet and tRNA Thr as depicted in **Supplemental Figure S13 B and D**. The secondary structures were optimized under these constraints with energy parameters re-scaled to 25°C and 0.1 M salt and isolated base-pairs allowed. The resulting minimum free energy structures were re-rendered using R2DT (McCann et al. 2025) to generate the visualizations shown in **Supplemental Fig. S13**.

## Data Availability

The direct RNA sequencing data is publicly accessible at the European Nucleotide Archive https://www.ebi.ac.uk/ena (accession number PRJEB85651).

## Acknowledgments

We thank Ute Kothe (University of Manitoba) for the generous gift of a Pus4 expression plasmid that we modified for use in this study. We thank Laura McKnight (Knight Campus for Accelerating Scientific Impact) for guidance on protein purification. We thank Marian Hettiaratchi, Parisa Hosseinzadeh and Justin Svendsen (Knight Campus for Accelerating Scientific Impact) for the use and training of their biolayer interferometry instrument. We thank Alice Barkan, Brad Nolen and Preeti Bhattacharjee (University of Oregon) for their comments on the manuscript.

## Contributions

D.M.G. and J.R.W. conceived the project. E.M.D., N.E.C.N., A.L.V., M.K., M.L.B., D.M.G. and J.R.W. conducted the investigation and performed the formal analysis, data curation and visualization. All authors contributed to the writing and editing of the manuscript.

## Ethics Declarations

The authors declare no competing interests

