## Supplemental Information for "Chronology of tRNA structural dynamics prior to and during interaction with a pseudouridine synthase"

A

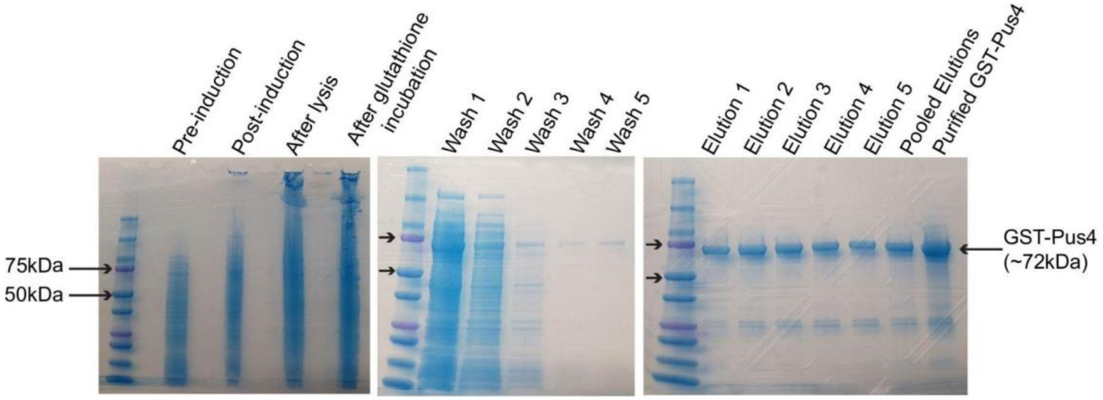

B

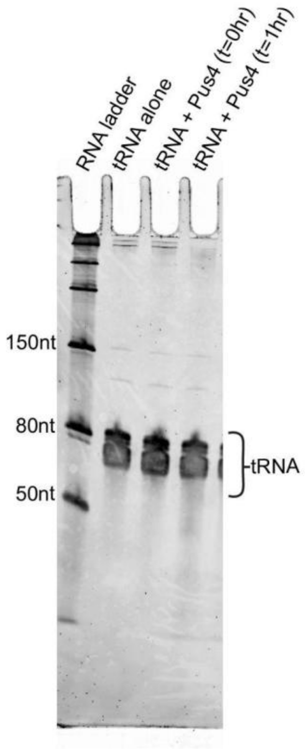

**SUPPLEMENTAL FIGURE S1.** Protein purification. (*A*) SDS-PAGE gels stained with GelCode Blue showing GST-Pus4 (72 kDa) present throughout purification. Upper and lower arrows indicate ladder standards of 75 and 50 kDa, respectively. (*B*) Urea PAGE gel showing the total yeast tRNA that remains without any appreciable degradation after incubation with purified Pus4 for 1 hour at room temperature.

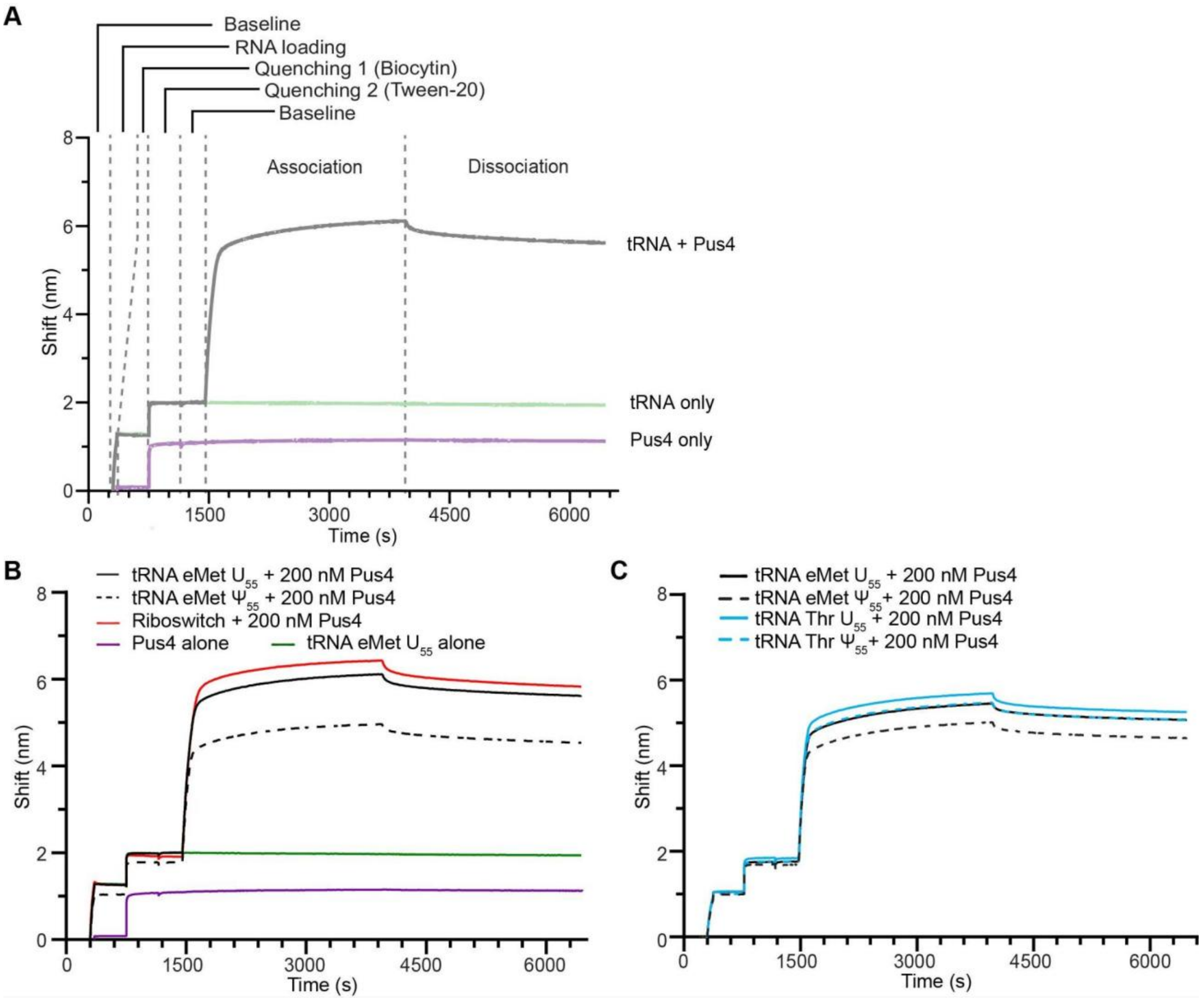

**SUPPLEMENTAL FIGURE S2.** Complete BLI traces of data from **Figure 2**. (*A*) Full BLI traces with each experimental step overlayed and labeled. Step (length of time): Baseline (300 s), RNA loading (~60 s), Quenching 1 (400 s), Quenching 2 (400 s), Baseline (300 s), Association (2500 s), Dissociation (2500 s). (*B*) Full BLI traces of 200 nM Pus4 showing association with $U_{55}$ (black solid) and $\Psi_{55}$ (black dashed) tRNA eMet, and $NAD^+$ riboswitch (red). The traces from the reference probes of Pus4 alone (purple) and $U_{55}$ tRNA eMet alone (green) are also shown. (*C*) Same as *B* but with $U_{55}$ tRNA eMet (black solid), $\Psi_{55}$ tRNA eMet (black dashed), $U_{55}$ tRNA Thr (blue solid), and $\Psi_{55}$ tRNA Thr (blue dashed).

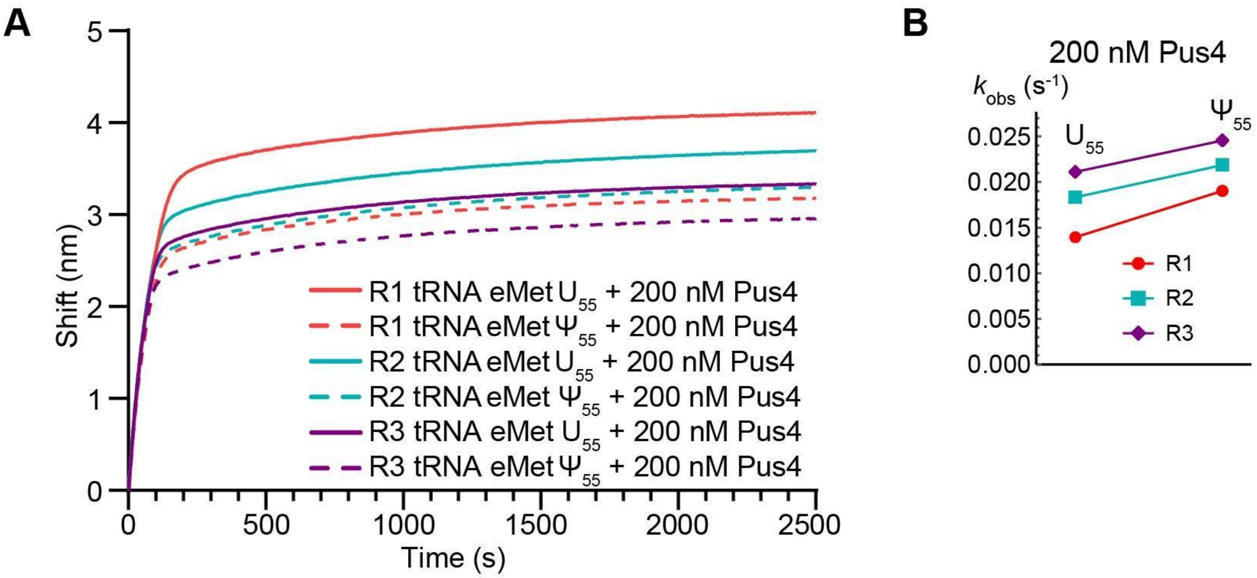

**SUPPLEMENTAL FIGURE S3.** BLI replicate data and resulting rates of fits to association phases of tRNA + Pus4 BLI curves. Each color represents a different BLI run. (*A*) Association phase of 200 nM Pus4 showing association with $U_{55}$ (solid) and $\Psi_{55}$ (dashed) tRNA eMet across three replicates (R1, R2, and R3). (*B*) $k_{obs}$ determined from a bi-exponential fit to the entire association phases of three runs (R1, R2 and R3) of tRNA eMet + 200 nM Pus4 on different days. While the rates themselves are variable between runs, $\Psi_{55}$ tRNA consistently exhibits a faster $k_{obs}$ than $U_{55}$ tRNA.

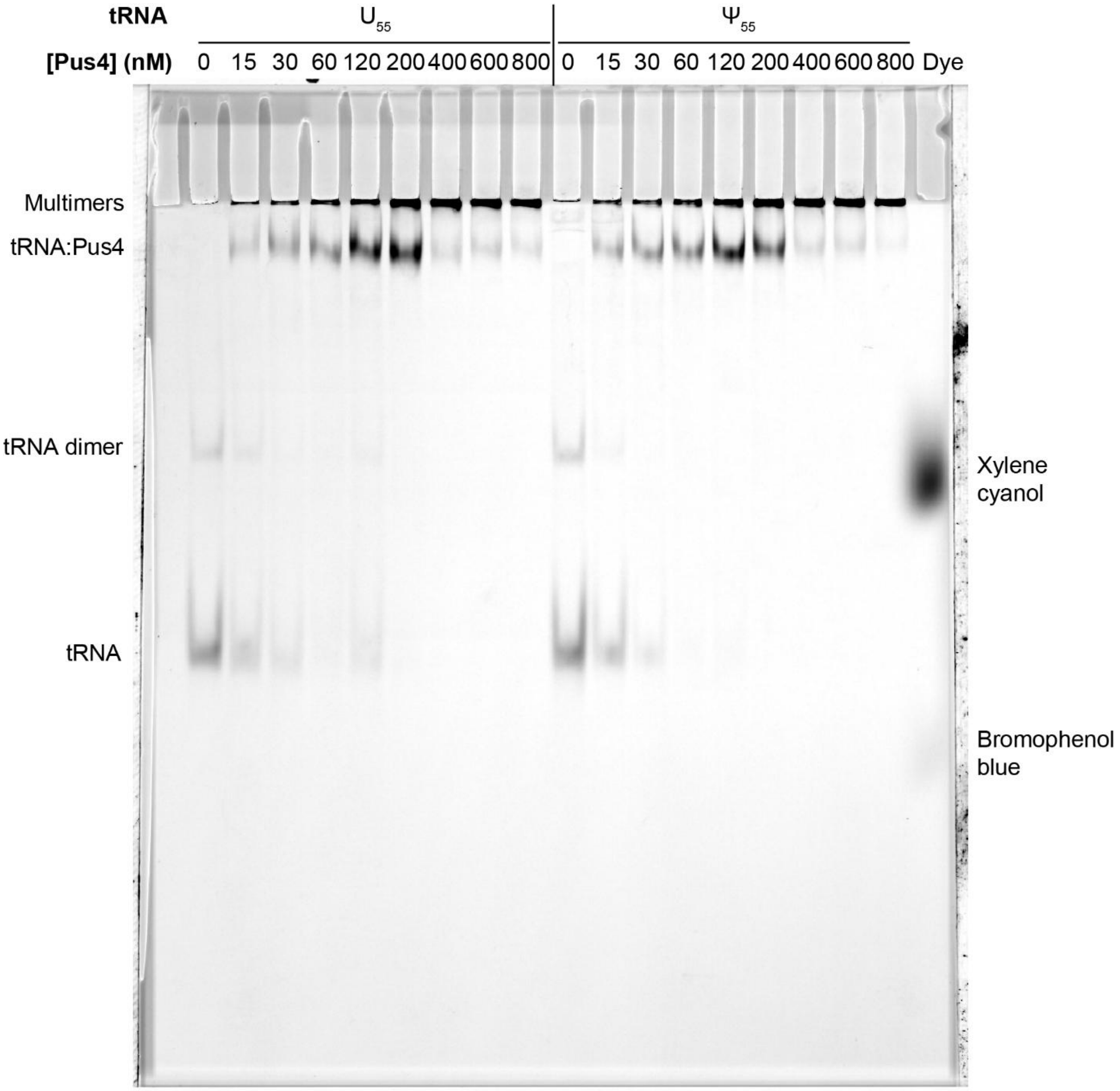

**SUPPLEMENTAL FIGURE S4.** Electrophoretic Mobility Shift Assay. $U_{55}$ or $\Psi_{55}$ tRNA eMet was incubated for 30 minutes in the presence of 0 to 800 nM Pus4 WT and the resulting protein-RNA assemblies were resolved on an 8% non-denaturing polyacrylamide gel. tRNA was visualized by detecting Cy5 fluorescence. The image was subjected to a 500-pixel rolling ball background subtraction. The slower-migrating band in the no-protein lanes may represent tRNA dimers. The far-right lane contains xylene cyanol and bromophenol blue as a reference.

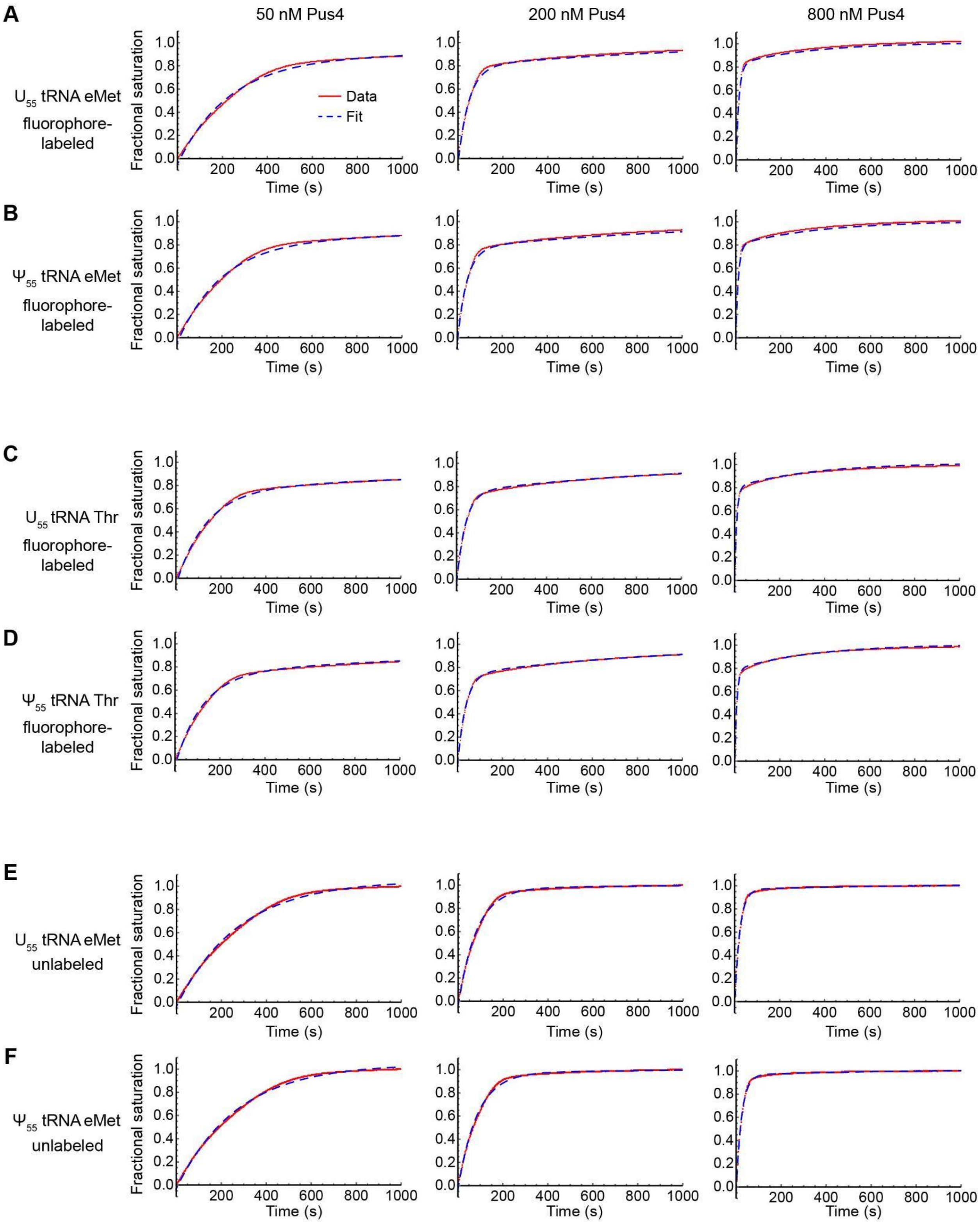

**SUPPLEMENTAL FIGURE S5.** Bi-exponential fits (blue dashed) to the association phases of BLI curves (red) at varying concentrations of Pus4. Left column: 50 nM; Middle: 200 nM; Right: 800 nM. (*A*) Fluorophore-labeled $U_{55}$ tRNA eMet. (*B*) Fluorophore-labeled $\Psi_{55}$ tRNA eMet. (*C*)

Fluorophore-labeled $U_{55}$ tRNA Thr. (*D*) Fluorophore-labeled $\Psi_{55}$ tRNA Thr. (*E*) Unlabeled $U_{55}$ tRNA eMet. (*F*) Unlabeled $\Psi_{55}$ tRNA eMet. The entire association phase was fitted for panels A-D, but the first 1000 s are displayed for ease of comparison.

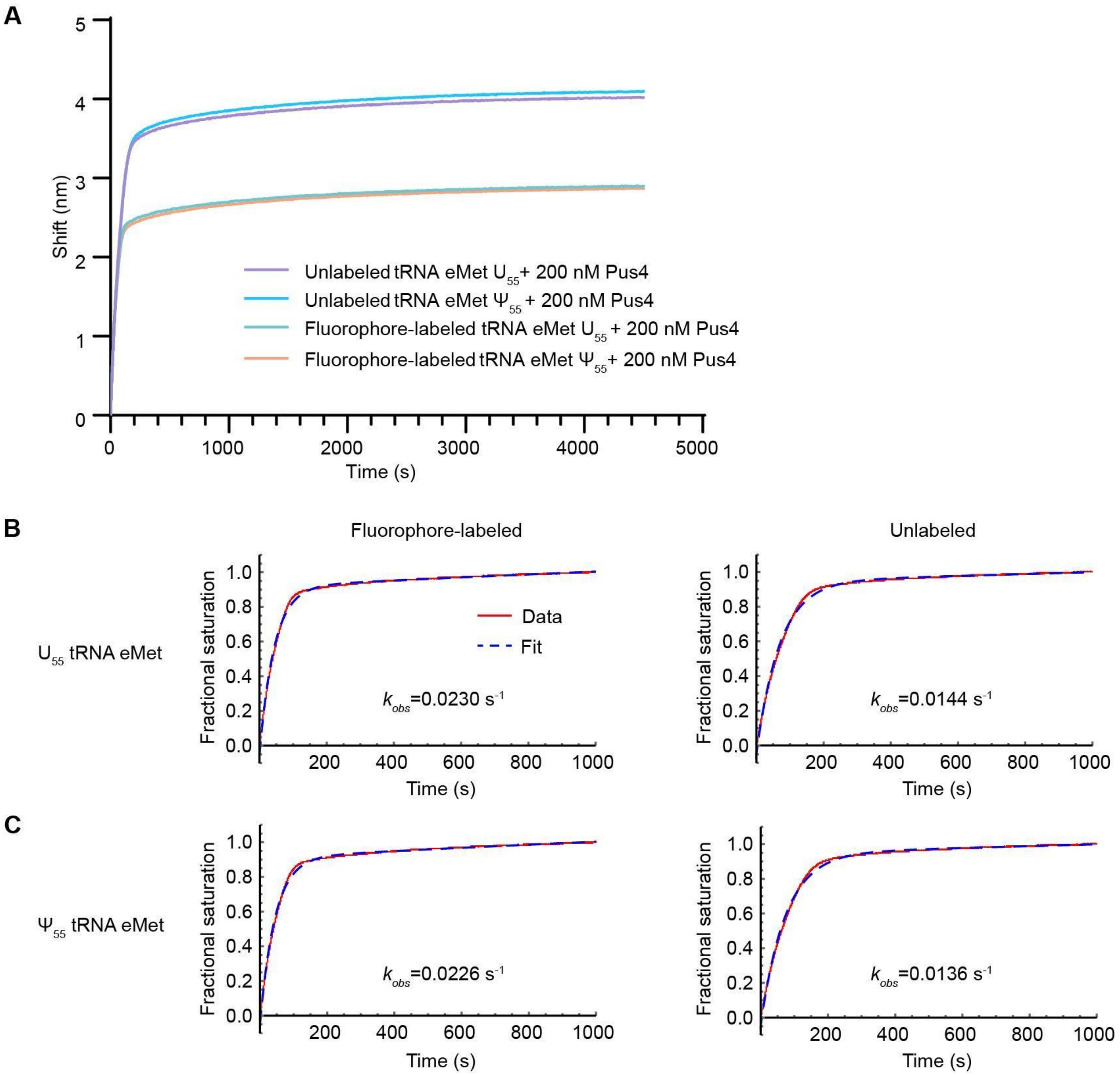

**SUPPLEMENTAL FIGURE S6:** Association phase of BLI curves and resulting fits of association to unlabeled and fluorophore-labeled tRNA eMet from within the same BLI run. (*A*) Association phase of 200 nM to unlabeled and fluorophore-labeled tRNA eMet. Traces are aligned to start at a shift of 0 nm. (*B*) Fits of the association phase of the BLI curve with $U_{55}$ tRNA eMet. (C) Fits of the association phase of the BLI curve with $\Psi_{55}$ tRNA eMet. The entire association phase was fitted, but the first 1000 s are displayed for ease of comparison. Pus4 shows a faster association rate to fluorophore-labeled tRNA eMet than unlabeled tRNA eMet, consistent with the comparison across BLI runs in **Fig. 4**.

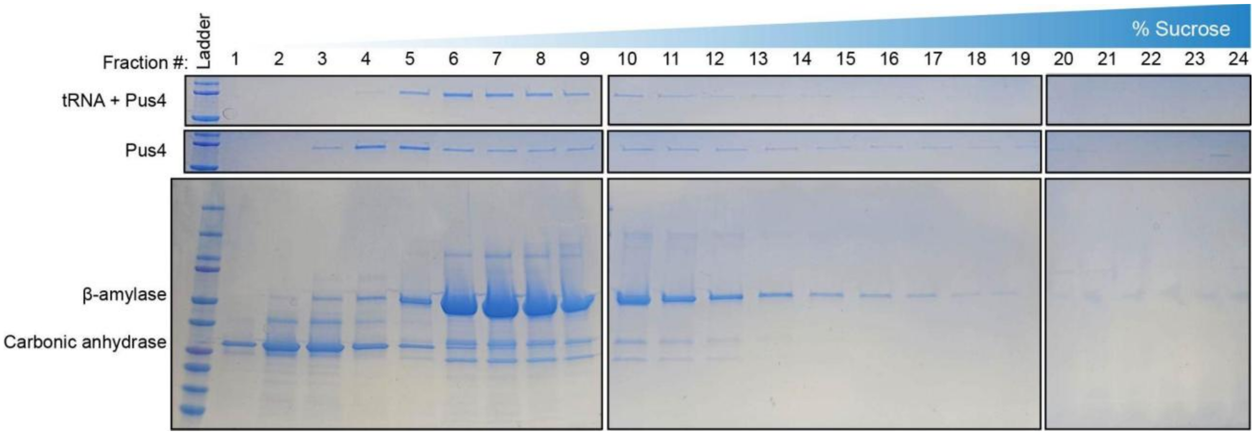

**SUPPLEMENTAL FIGURE S7.** Sucrose gradient ultracentrifugation. SDS-PAGE gels stained with GelCode Blue showing GST-Pus4 (72 kDa) with and without incubation with yeast tRNA as separated by a sucrose gradient (10–30% sucrose). Fraction 1 (low % sucrose) contains smaller complexes or monomers, while fraction 24 (high % sucrose) contains larger complexes. A control gradient was run in parallel containing purified β-amylase and carbonic anhydrase, which should form native protein complexes of 200 kDa (monomer is 50 kDa) and 26 kDa, respectively.

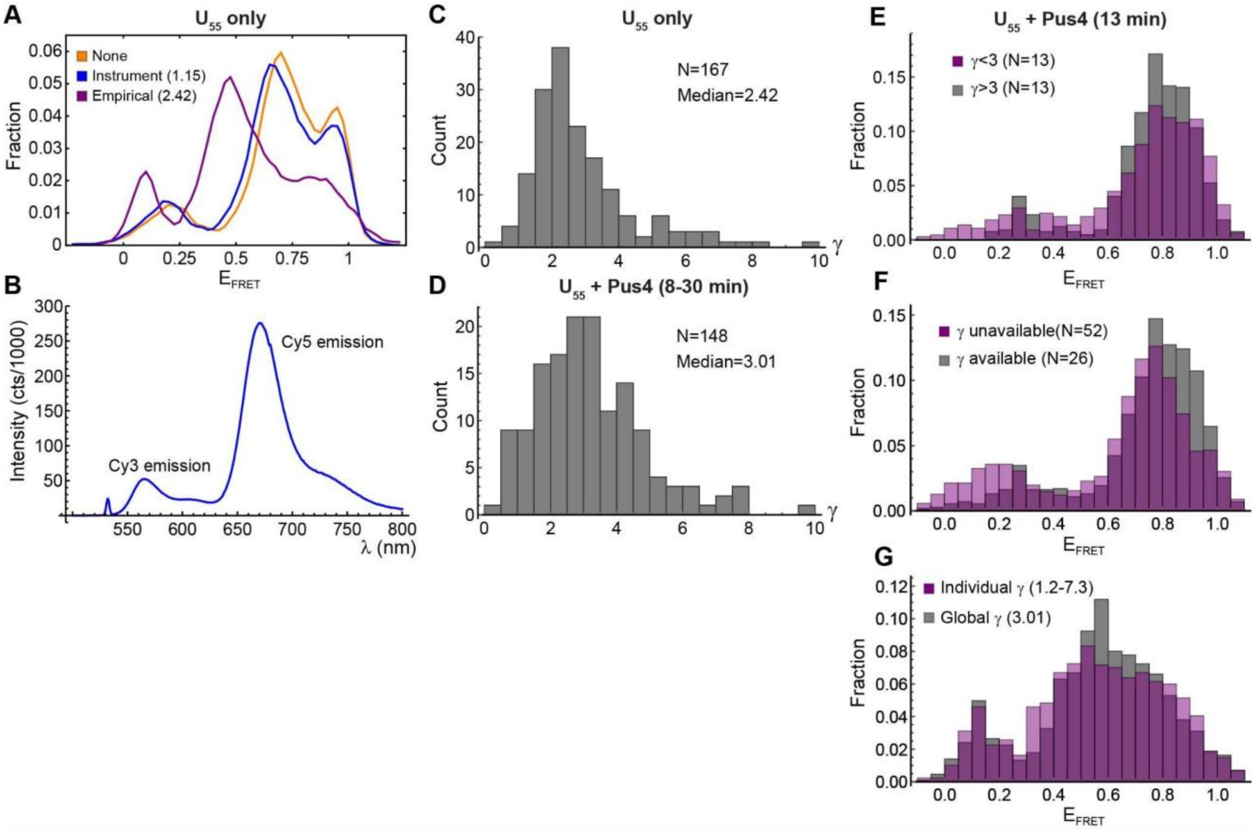

**SUPPLEMENTAL FIGURE S8.** Gamma factor correction. (*A*) Comparison of $U_{55}$ tRNA eMet $E_{FRET}$ histograms compiled from traces without γ factor correction (orange), traces corrected with a value of γ=1.15 calculated from the properties of the TIRF microscopy instrument's emission path optics and camera (blue) or traces corrected with empirically determined median value of γ=2.42 (purple, see panel *C*). (*B*) Bulk emission spectrum of $U_{55}$ tRNA under excitation at 532 nm. Much brighter Cy5 than Cy3 emission indicates that the skew toward high apparent $E_{FRET}$ is not instrument-specific. (*C,D*) Histograms of γ factor values obtained from $U_{55}$ tRNA eMet traces recorded in the absence (*C*) and presence (*D*) of Pus4 (details in **Methods**). "N" indicates the number of traces included in each histogram. (*E-G*) $E_{FRET}$ histograms justifying the use of a global γ factor correction (data on $U_{55}$ tRNA eMet+Pus4). (*E*) Uncorrected traces with γ above the median (gray) have narrower features but a similar overall shape to traces with γ below the median. (*F*) Uncorrected traces for which γ could be calculated (gray) skew toward higher $E_{FRET}$ than traces for which it could not be calculated, suggesting that limiting the analysis to molecules for which an individual γ can be calculated (those in which the acceptor fluorophore photobleaches before the donor) would distort the results. (*G*) Traces (N=26) corrected with individual values of γ yield a broader but otherwise similar histogram to traces corrected with the global value of γ=3.01.

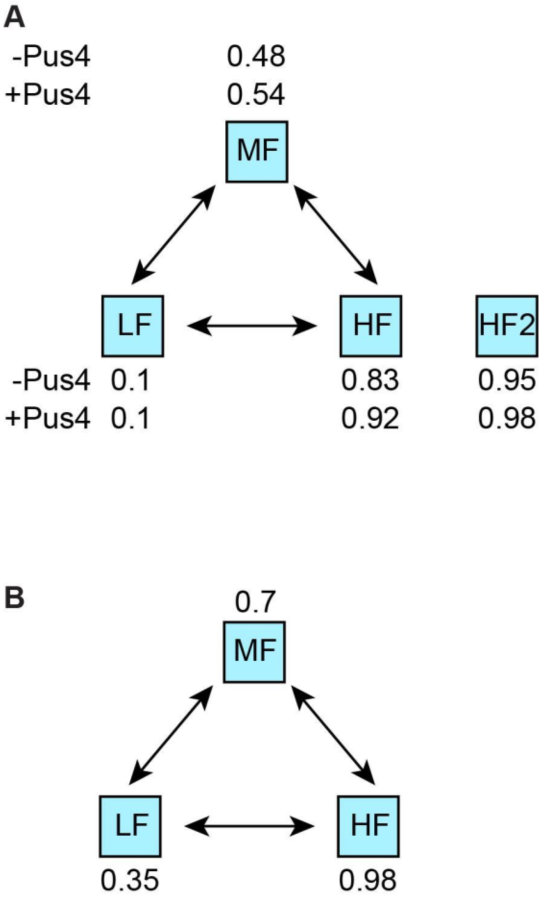

**SUPPLEMENTAL FIGURE S9.** Graphical depictions of the models used for hidden Markov modeling of (*A*) tRNA eMet and (*B*) tRNA Thr. The starting guess for the FRET efficiency of each state is indicated. HMM of tRNA eMet in the presence of Pus4 utilized the same model with different starting guesses (second line of numbers in panel *A*).

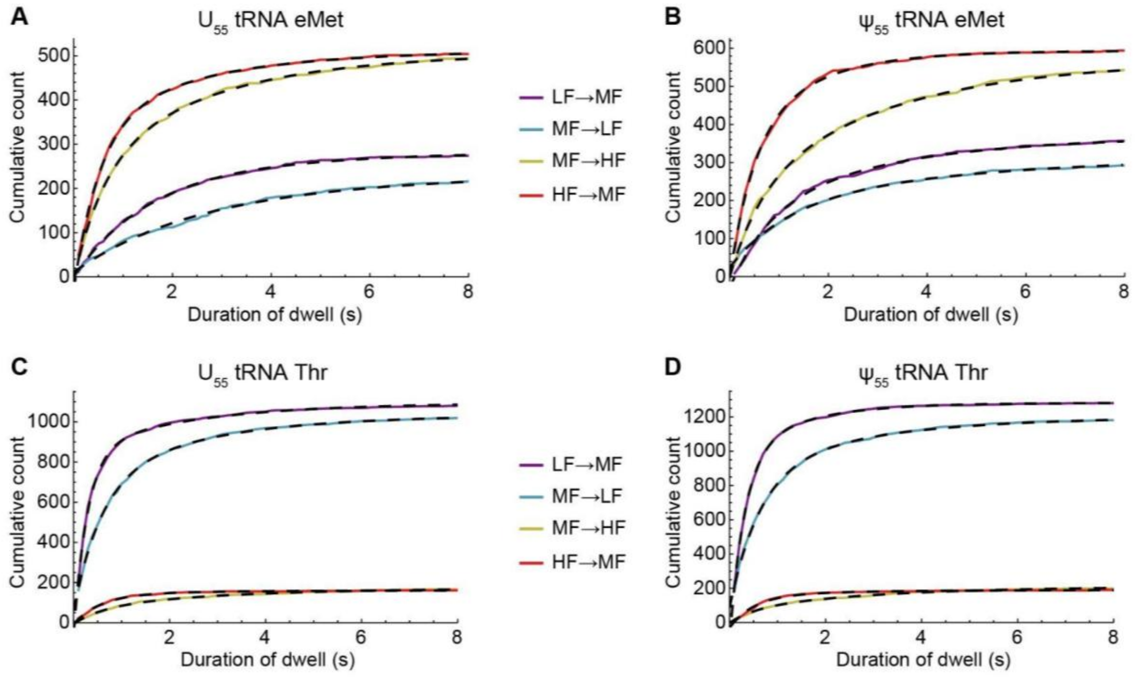

**SUPPLEMENTAL FIGURE S10.** Dwell times of tRNAs in each conformational state in the absence of Pus4. (*A*) Number of times $U_{55}$ tRNA eMet dwelled in a state for a duration less than or equal to the time indicated on the x-axis. Purple: dwell times in the LF state prior to transitioning to the MF state. Blue: dwell times in MF prior to transitioning to LF. Yellow: dwell times in MF prior to transitioning to HF. Red: Dwell times in HF prior to transitioning to MF. Bi-exponential fits are overlaid in dashed black lines. (*B*) Same as *A* but for $\Psi_{55}$ tRNA eMet. (*C*) Same as *A* but for $U_{55}$ tRNA Thr. (*D*) Same as *A* but for $\Psi_{55}$ tRNA Thr.

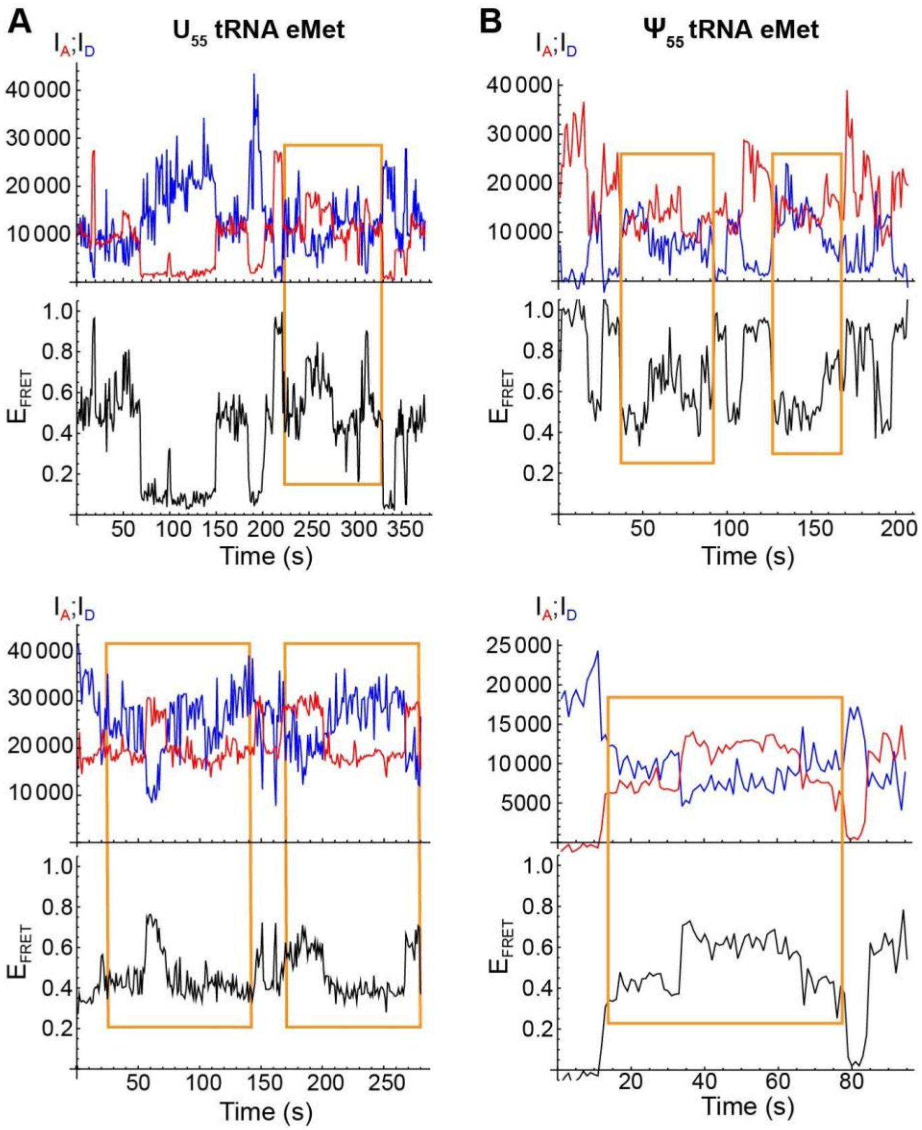

**SUPPLEMENTAL FIGURE S11.** Example traces showing interconversion between MF microstates (highlighted by orange boxes) in the presence of Pus4 for (*A*) $U_{55}$ tRNA eMet or (*B*) $\Psi_{55}$ tRNA eMet.

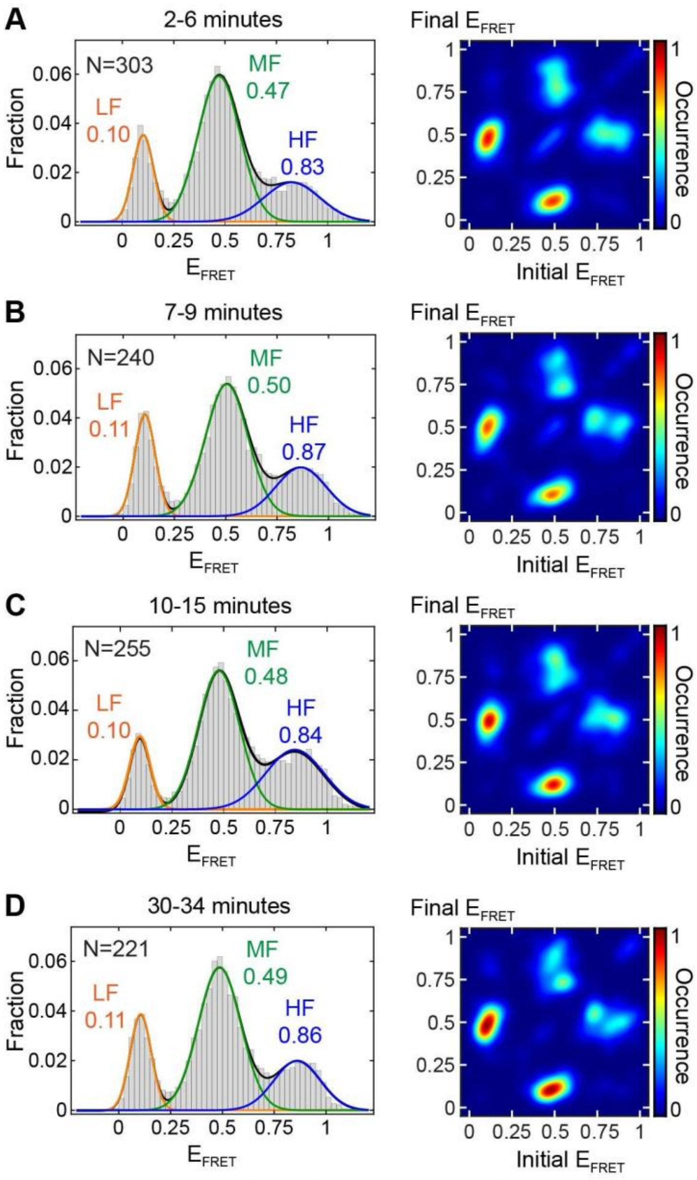

**SUPPLEMENTAL FIGURE S12.** $U_{55}$ tRNA eMet smFRET timecourse without the addition of Pus4. Histograms (left column) and TODPs (right column) recorded (*A*) 2–6 minutes after introduction of imaging buffer; (*B*) 7–9 minutes; (*C*) 10–15 minutes; (*D*) 30–34 minutes.

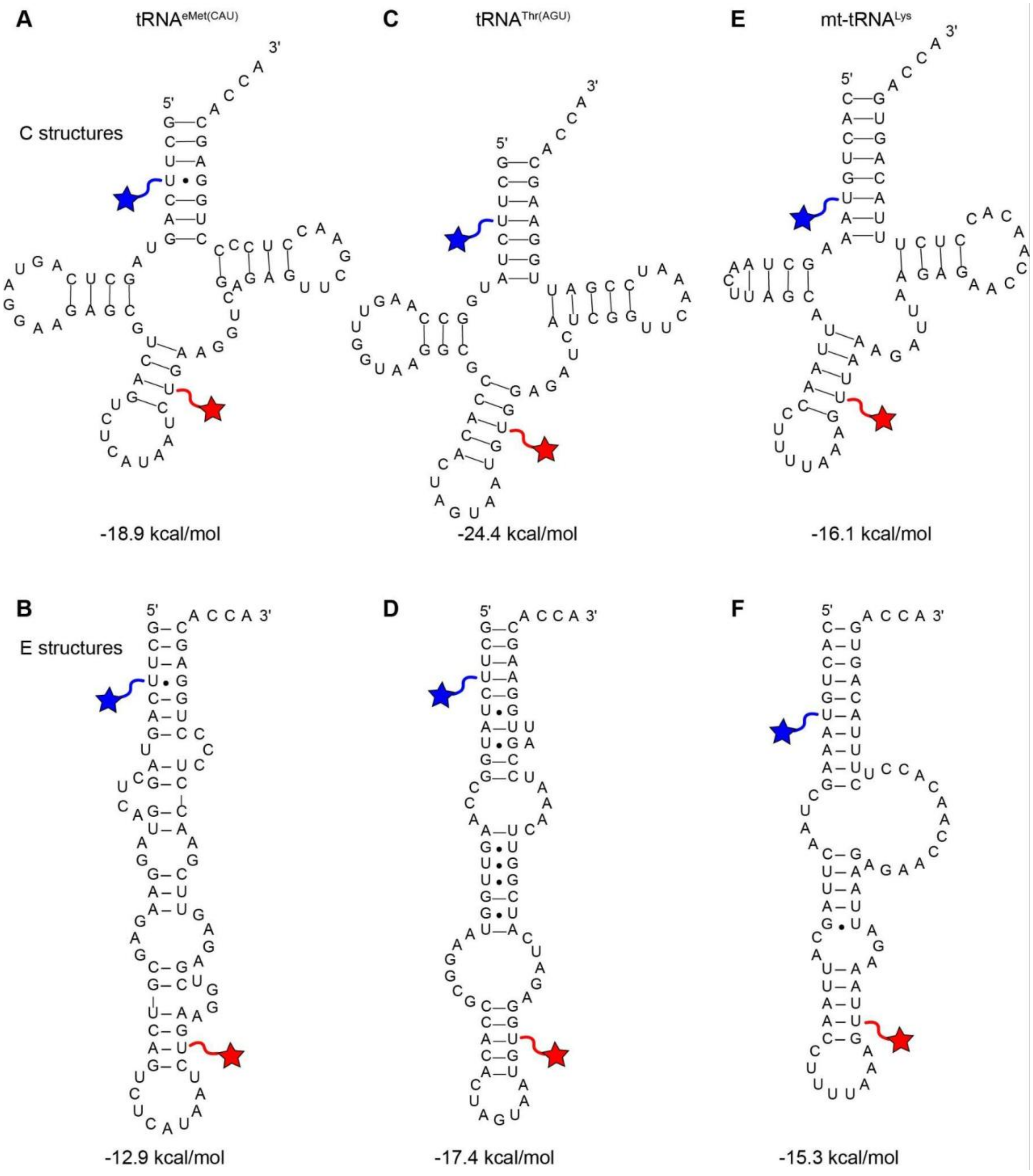

**SUPPLEMENTAL FIGURE S13.** Secondary structures and associated free energies at 25°C and 0.1 M salt predicted using the RNAfold web server (see **Methods** for details). (*A*) Predicted C structure of tRNA eMet. (*B*) Plausible E-like structure of tRNA eMet. (*C*) Predicted C structure of tRNA Thr. (*D*) Plausible E-like structure of tRNA Thr. (*E*) C structure of human mitochondrial $tRNA^{Lys}$ from Voigts-Hoffmann et al. 2007. (*F*) E structure of human mitochondrial $tRNA^{Lys}$ from Voigts-Hoffmann et al. 2007.

**SUPPLEMENTAL TABLE S1.** Yeast strains, plasmids and plasmid insert sequences.

**Yeast strains**

| **Strain** | **Description** | **From** |
|---|---|---|
| YDG1 | wild-type, BY4741 | Horizon/Dharmacon Reagents |
| YDG127 | *pus4*Δ::KanMX | Yeast Deletion Collection (Giaever et al. 2002) |

**Plasmids**

| **Construct** | **Description** | **From** |
|---|---|---|
| PDG322 | To express WT Pus4 in *E. coli*; pGEX-5X-3-ScPus4 | This study, modified from plasmid gifted by Dr. Ute Kothe |

**Insert sequences**

Sequence of expressed wild-type Pus4 protein (including GST tag, Factor Xa cleavage site, and Pus4 protein sequence):

ATGTCCCCTATACTAGGTTATTGGAAAATTAAGGGCCTTGTGCAACCCACTCGACTTCTTTT
GGAATATCTTGAAGAAAAATATGAAGAGCATTTGTATGAGCGCGATGAAGGTGATAAATGG
CGAAACAAAAAGTTTGAATTGGGTTTGGAGTTTCCCAATCTTCCTTATTATATTGATGGTGA
TGTTAAATTAACACAGTCTATGGCCATCATACGTTATATAGCTGACAAGCACAACATGTTGG
GTGGTTGTCCAAAAGAGCGTGCAGAGATTTCAATGCTTGAAGGAGCGGTTTTGGATATTAG
ATACGGTGTTTCGAGAATTGCATATAGTAAAGACTTTGAAACTCTCAAAGTTGATTTTCTTAG
CAAGCTACCTGAAATGCTGAAAATGTTCGAAGATCGTTTATGTCATAAAACATATTTAAATG
GTGATCATGTAACCCATCCTGACTTCATGTTGTATGACGCTCTTGATGTTGTTTTATACATG
GACCCAATGTGCCTGGATGCGTTCCCAAAATTAGTTTGTTTTAAAAAACGTATTGAAGCTAT
CCCACAAATTGATAAGTACTTGAAATCCAGCAAGTATATAGCATGGCCTTTGCAGGGCTGG
CAAGCCACGTTTGGTGGTGGCGACCATCCTCCAAAATCGGATCTGATCGAAGGTCGTGGG
ATCCCCAGGAATTCGGGGAACATGAATGGAATATTTGCTATTGAGAAGCCTAGCGGTATTA
CATCCAATCAGTTTATGCTAAAGTTGCAACATGCTTTAACTAAAAGTCAAGTATTCTCGAAG
GAAATTCAAAGGGCAACAGCGGAAAGAAAGCAGCAATATGAAAAACAGACAGGTAAGAAG
GCCAGCAAGAGGAAACTTCGTAAAGTCTCAAAGGTAAAGATGGGACACGGAGGCACTTTG
GATCCGTTGGCCTCTGGTGTCCTGGTAATTGGGATTGGAGCGGGCACCAAAAAACTTGCA
AATTATCTTTCAGGCACCGTTAAAGTCTATGAAAGTGAAGCCCTGTTTGGTGTTTCCACTAC
TTCAGGAGATGTAGAAGGTGAGATTTTGTCCCAGAACTCCGTTAAACATTTAAATTTCGATG
ATTTAAAGACTGTGGAGGAAAAATTTGTTGGACAATTAAAGCAGACGCCGCCAATATACGC
AGCTTTGAAGATGGATGGCAAACCCCTGCATGAATATGCTCGAGAAGGAAAGCCTTTACCT
AGAGCCATTGAGCCCAGACAAGTTACTATTTATGACTTGAAGGTGTTTTCAGACTCTCTTAA
ACGTGATCATGACTATCCCTTATTGAGGCCCACTACAGAAGAAGCTGTTGATACCGTGAAA
AATTTGAATGCAAACATGCTAAACGATGTTTTATACTTTTCAAAAGAATATACAGAGAAGCAC

GGCCTTGATTCTGAGGTGGCAAAGGTAGAAGAACCATTTCCGCTGAGTGAACAAGAAGAG
CAAGAAATCCAAAAAGAAGGAGATTCTTATAGGGCTCCAAAATTGCACTTCAAAGCTAATGT
GTCTTCTGGAACATACATAAGGTCCTTAGTGAGCGACATCGGTAAATCTATGAGGAGCTCA
TGTTATATGGTGAAATTGATACGTTTGCAACAGCAGGACTGGTCTCTTGAAAAAAATAATGT
GTTCCAATTAACGGATTTTACTGAAAGAGACGAGAAGGTGTGGAGTAAGGTACTGGAGAAG
GTTCTAGACGAGGGAGCAACTGTTGATGTTATAGAAGAGCTAAAGAAAGCAGAAAAGGAAA
TACCAGCGGACGTGAAGGAATGTATAGTTTCGAGTGATCAACCTGGTGATGAGGCTACGG
CTGAAACAATTGAAACTGCTAACGCCGAAGAACATTCTAATACACTAAAAAGAAAAATCGAA
CAGGTGTAA

**SUPPLEMENTAL TABLE S2.** Fluorophore intensities.

The mean intensity across all frames of all donor or acceptor traces within each analyzed movie was determined. The table reports the mean and standard deviation of this value across the indicated number of movies. Time ranges indicate the time window following injection of imaging buffer containing OSS either with or without Pus4 during which all movies in a condition were recorded. Measurements are on tRNA eMet unless otherwise noted.

| Condition | Donor intensity | Acceptor intensity | Movies |
|---|---|---|---|
| $U_{55}$ only | 2500±150 | 3300±340 | 3 |
| $U_{55}$ + Pus4 2–6 min. | 9400±1670 | 9200±970 | 5 |
| $U_{55}$ + Pus4 7–9 min. | 10000±2760 | 10000±1290 | 4 |
| $U_{55}$ + Pus4 10–15 min. | 12700±950 | 11000±1260 | 4 |
| $U_{55}$ + Pus4 30–34 min. | 11000±1080 | 11000±1070 | 3 |
| $\Psi_{55}$ only | 2500±480 | 2800±400 | 3 |
| $\Psi_{55}$ + Pus4 2–6 min. | 10800±500 | 10000±1720 | 3 |
| $\Psi_{55}$ + Pus4 7–9 min. | 10700±800 | 12100±620 | 2 |
| $\Psi_{55}$ + Pus4 10–15 min. | 11300±930 | 11700±110 | 3 |
| $\Psi_{55}$ + Pus4 30–34 min. | 12000±1620 | 11400±850 | 3 |
| $U_{55}$ alone 2–6 min.* | 4600±280 | 4000±620 | 2 |
| $U_{55}$ alone 7–9 min.* | 4270 | 4430 | 1 |
| $U_{55}$ alone 10–15 min.* | 4130±280 | 4320±270 | 2 |

| | | | |
|---|---|---|---|
| $U_{55}$ alone 30–34 min.* | 4500±520 | 3930±30 | 2 |
| $\Psi_{55}$ alone 10–15 min.* | 4320±50 | 4100±80 | 2 |
| $U_{55}$ tRNA$^{Thr}$ 10–15 min.* | 3200±100 | 5100±160 | 2 |
| $\Psi_{55}$ tRNA$^{Thr}$ 10–15 min.* | 3400±340 | 5500±230 | 3 |

*Recorded with an electron multiplication (EM) gain setting of 450. All others recorded with a setting of 300.

**SUPPLEMENTAL TABLE S3.** Oligonucleotides

| **Description** | **Sequence (5′ to 3′)** | **Supplier and Purification** | **Nucleic Acid** |
|---|---|---|---|
| eMet CAU 5′ oligo | biotin-GCUU[Cy3]CAGUAGCUCAGUAGGAAGAGCGUCAGUCUCAUAAU | Dharmacon, HPLC | RNA |
| eMet CAU 3′ oligo | phosphate-CU[Cy5]GAAGGUCGAGAGUUCGAACCUCCCCUGGAGCACCA | Dharmacon, HPLC | RNA |
| eMet CAU 3′ with Ψ | phosphate-CU[Cy5]GAAGGUCGAGAGU**Ψ**CGAACCUCCCCUGGAGCACCA | Dharmacon, HPLC | RNA |
| Thr AGU 5′ oligo | biotin-GCUU[Cy3]CUAUGGCCAAGUUGGUAAGGCGCCACACUAGUAAU | Dharmacon, HPLC | RNA |
| Thr AGU 3′ oligo | phosphate-GU[Cy5]GGAGAUCAUCGGUUCAAAUCCGAUUGGAAGCACCA | Dharmacon, HPLC | RNA |
| Thr AGU 3′ with Ψ | phosphate-GU[Cy5]GGAGAUCAUCGGU**Ψ**CAAAUCCGAUUGGAAGCACCA | Dharmacon, HPLC | RNA |
| Splint for 5′ and 3′ portions of tRNA eMet | GGTGGGTCTGCTTTGGTGCTCCAGGGGAGGTTCGAACTCTCGACCTTCAGATTATGAGACTGACGCTCTTCCTACTGAGCTACTGAAGCATTTTGGGTAC | IDT, PAGE | DNA |

| | | | |
|---|---|---|---|
| Splint for 5′ and 3′ portions of tRNA Thr | GGTGGGTCTGCTTTGGTGCTTCCAATCGGAT TTGAACCGATGATCTCCACATTACTAGTGTG GCGCCTTACCAACTTGGCCATAGAAGCATTT TGGGTAC | IDT, PAGE | DNA |
| NAD$^+$ riboswitch | biotin-AUCUAUAGAGCGUU(Cy5)GCGUCCGAAAGU CUAAACAGACACGGCUCUUUAAAAACAAAA GGAGA(Cy3) | Dharmacon, HPLC | RNA |
| eMet CAU, no fluorophores | biotin-GCUUCAGUAGCUCAGUAGGAAGAGCGUCA GUCUCAUAAUCUGAAGGUCGAGAGUUCGAA CCUCCCCUGGAGCACCA | Dharmacon, HPLC | RNA |
| eMet CAU with Ψ, no fluorophores | biotin-GCUUCAGUAGCUCAGUAGGAAGAGCGUCA GUCUCAUAAUCUGAAGGUCGAGAGU**Ψ**CGA ACCUCCCCUGGAGCACCA | Dharmacon, HPLC | RNA |

**SUPPLEMENTAL TABLE S4.** Direct RNA Sequencing Data Files

**Sample** is the name of the sample as referred to in Dennis and Conoan Nieves *et al*. 2025.

**Fast5 and FastQ file** are the names of the files containing the basecalled sequencing data for the specified sample. Files from this study are available at the European Nucleotide Archive (https://www.ebi.ac.uk/ena) with study accession number PRJEB85651. Files from Shaw et al. 2024 reanalyzed in this study are available with study accession number PRJEB68201.

| Accession | Sample | Fast5 file | FastQ file |
|---|---|---|---|
| PRJEB85651 | WT Pus4 *in vitro* | 20240718_RNA002_UO_sCer_tRNA_pus4deletion_in_vitro_WT_pus4.tar.gz | 20240718_RNA002_UO_sCer_tRNA_pus4deletion_in_vitro_WT_pus4_noU.fastq.gz |
| PRJEB85651 | WT Pus4 no $MgCl_2$ *in vitro* | 20240813_RNA002_UO_sCer_tRNA_pus4deletion_in_vitro_WT_pus4_noMgCl.tar.gz | 20240813_RNA002_UO_sCer_tRNA_pus4deletion_in_vitro_WT_pus4_noMgCl_noU.fastq.gz |
| PRJEB68201 | *in vivo* WT replicate 1 | WT_tRNAs_rep1_7_22_21_fast5s.tar.gz | WT_tRNAs_rep1_7_22_noU.fastq.gz |
| PRJEB68201 | *in vivo* WT replicate 2 | WT_tRNAs_rep2_8_6_21_fast5s.tar.gz | WT_tRNAs_rep2_8_6_21_noU.fastq.gz |
| PRJEB68201 | *in vivo* WT | WT_tRNAs_rep3_9_17_21 | WT_tRNAs_rep3_9_17_21_noU.f |

| | | | |
|---|---|---|---|
| | replicate 3 | _fast5s.tar.gz | astq.gz |
| PRJEB68201 | *in vivo* *pus4*Δ replicate 1 | pus4_tRNAs_rep1_06_30_21_fast5s.tar.gz | pus4_tRNAs_rep1_06_30_21_noU.fastq.gz |
| PRJEB68201 | *in vivo* *pus4*Δ replicate 2 | pus4_tRNAs_rep2_07_19_21_fast5s.tar.gz | pus4_tRNAs_rep2_07_19_21_noU.fastq.gz |
| PRJEB68201 | *in vivo* *pus4*Δ replicate 3 | pus4_tRNAs_rep3_10_26_21_fast5s.tar.gz | pus4_tRNAs_rep3_10_26_21_noU.fastq.gz |
